# An Excitatory Projection from the Basal Forebrain to the Ventral Tegmental Area Underlying Anorexia-like Phenotypes

**DOI:** 10.1101/2023.05.05.539422

**Authors:** Jing Cai, Yuanzhong Xu, Zhiying Jiang, Yanyan Jiang, Claire Young, Hongli Li, Joshua Otiz-Guzman, Yizhou Zhuo, Yulong Li, Yong Xu, Benjamin R. Arenkiel, Qingchun Tong

**Author notes:** Correspondence authors: Ben Arenkiel, PhD, or Qingchun Tong, PhD.

## Abstract

Adaptation to potential threat cues in feeding regulation is key for animal survival. Maladaptation in balancing internal energy needs and external threat cues may result in eating disorders. However, brain mechanisms underlying such maladaptation remain elusive. Here, we identified that the basal forebrain (BF) sends glutamatergic projections to glutamatergic neurons in the ventral tegmental area (VTA). Glutamatergic neurons in both the BF and the VTA displayed correlated responses to various external stressors. Notably, *in vivo* manipulation of BF terminals in the VTA revealed that the glutamatergic BF➔VTA circuit reduces appetite, increases locomotion, and elicits avoidance. In consistence, activation of VTA glutamatergic neurons reduced body weight, blunted food motivation, and caused hyperactivity with behavioral signs of anxiety, all hallmarks of typical anorexia phenotypes. Importantly, activation of BF glutamatergic terminals in the VTA reduced dopamine release in the nucleus accumbens (NAc). Collectively, our results point to overactivation of the glutamatergic BF➔VTA circuit as a potential cause of anorexia-like phenotypes involving reduced dopamine release.

## Introduction

With ever-changing environmental threat cues and internal nutritional checkpoints, an appropriate decision to engage in feeding behaviors is key for animal survival. Ongoing physical and perceived environmental disturbances can override internal energy needs and suppress feeding behaviors^1,2^. Maladaptation in balancing internal energy needs and external threat cues may result in anorexia or overeating, associated with debilitating malnutrition or obesity. In humans, heightened stress responses triggered by external cues reduce appetite and food motivation^3,4^. Specifically, in patients diagnosed with anorexia nervosa, an eating disorder with self-imposed starvation that has the highest mortality rate among mental disorders, there is a high prevalence of generalized anxiety disorders^5,6^, suggesting a profound impact of heightened stress responses on feeding behaviors. However, the brain circuits involved in sensing external stress cues to modulate feeding behaviors appropriately remain elusive.

In addition to its well-known functions in regulating arousal, wakefulness, learning, and memory, the basal forebrain (BF) has also been shown to sense environmental cues and process sensory input^7–12^. The BF contains cholinergic, gamma-aminobutyric acid (GABAergic), and glutamatergic (glu+) neurons, and all these neuron groups respond to external stimuli^9–11^. Interestingly, recent results also suggest the importance of BF neurons in feeding regulation. BF GABAergic neurons promote food consumption and drive high-calorie food intake^13,14^. Loss of BF cholinergic neurons induces massive obesity due to increased food intake, while chronic activation of BF^glu+^ neurons leads to reduced food intake and starvation^15,16^. Notably, BF^glu+^ neurons respond to various sensory inputs, including odors with different valences, predator cues, and other physical threats associated with reduced feeding in animals^15,17,18^. These observations posit that the BF serves as a functional hub for sensing environmental cues, thus capable of profoundly impacting feeding behaviors.

The ventral tegmental area (VTA) is a well-studied dopamine (DA)-enriched midbrain structure involved in reward responses, motivation, learning, and memory ^19–21^. Consistent with its role in reward consumption, the VTA has also been shown to modulate feeding, especially with respect to hedonic behaviors^22^. In particular, DA release from the VTA signals for high-fat diet (HFD) intake, and coincides with HFD-induced activity changes in hypothalamic agouti-related protein (AgRP) neurons, which have been established as regulatory feeding neurons^22–24^. In addition to DA neurons, the VTA consists of GABAergic and glu+ neurons, which have been shown to gate DA signals^25–29^. VTA GABAergic neurons form local connections with DA neurons to inhibit DA release and reduce reward consumption ^28–30^. Similarly, VTA^glu+^ neurons send direct projections to various brain regions as well as local DA neurons^25,27^, and play a complicated role in reward processing^31–34^. Yet, how VTA neurons integrate environmental cues to regulate feeding behaviors is unclear.

Here we show that BF^glu+^ neurons send direct projections to the VTA, preferentially targeting glu+ neurons. Both BF^glu+^ and VTA^glu+^ neurons responded to various environmental stimuli in a correlative manner. *In vivo* activation of the BF^glu+^➔VTA circuit inhibited feeding and led to behavioral avoidance. Activation of downstream VTA^glu+^ neurons led to hypophagia, diminished food motivation, hyperactivity, behavioral signs of anxiety, and body weight reduction, all resembling typical symptoms of anorexia. Furthermore, activation of this glutamatergic BF➔VTA circuit was associated with reduced DA release in the nucleus accumbens (NAc), indicating a potential involvement of DA release in some of the observed anorexia-like phenotypes.

## Results

### The BF sends glutamatergic projections to VTA^glu+^ neurons

To identify the downstream targets of BF^glu+^ neurons, we conducted channelrhodopsin-2 (ChR2) –assisted circuit mapping. Given the preferential localization of ChR2 on cell membrane, ChR2 fused with a fluorescent reporter can be used to trace downstream projection sites^35^. To specifically target BF^glu+^ neurons, we used Vglut2-ires-Cre, a mouse strain that expresses Cre from the vesicular glutamate transporter 2 (Vglut2, SLC17a6) locus^36^, and stereotaxically delivered conditional AAV5-EF1a-DIO-ChR2-EYFP viral particles to the BF (Figures 1a and 1b). Consistent with previous studies^15,17^, ChR2-EYFP positive fibers were detected in the lateral habenula (LHb), the lateral hypothalamus (LH), and other brain regions (Figure S1b). Of note, we also found abundant EYFP positive fibers in the VTA (Figure 1c). To verify the projection pattern, we performed retrograde adeno-associated viral (AAVrg)-based tracing experiments by injecting AAVrg-Ef1a-mCherry-IRES-Flp into the VTA, and AAV-DJ8-hSyn-Con/Fon-EYFP into the BF of Vglut2-ires-Cre mice (Figures 1d-f). Given that the AAV-DJ8-hSyn-Con/Fon-EYFP vector expresses EYFP in a Cre and Flp co-dependent manner, in this configuration, EYFP selectively marked BF^glu+^ neurons that project to the VTA (Figure 1f). We found that EYFP-positive neuronal fibers were also present in the LHb in addition to the VTA (Figures 1g and 1h), suggesting that these VTA-projecting BF^glu+^ neurons send collateral projections to the LHb.

**Figure 1:**
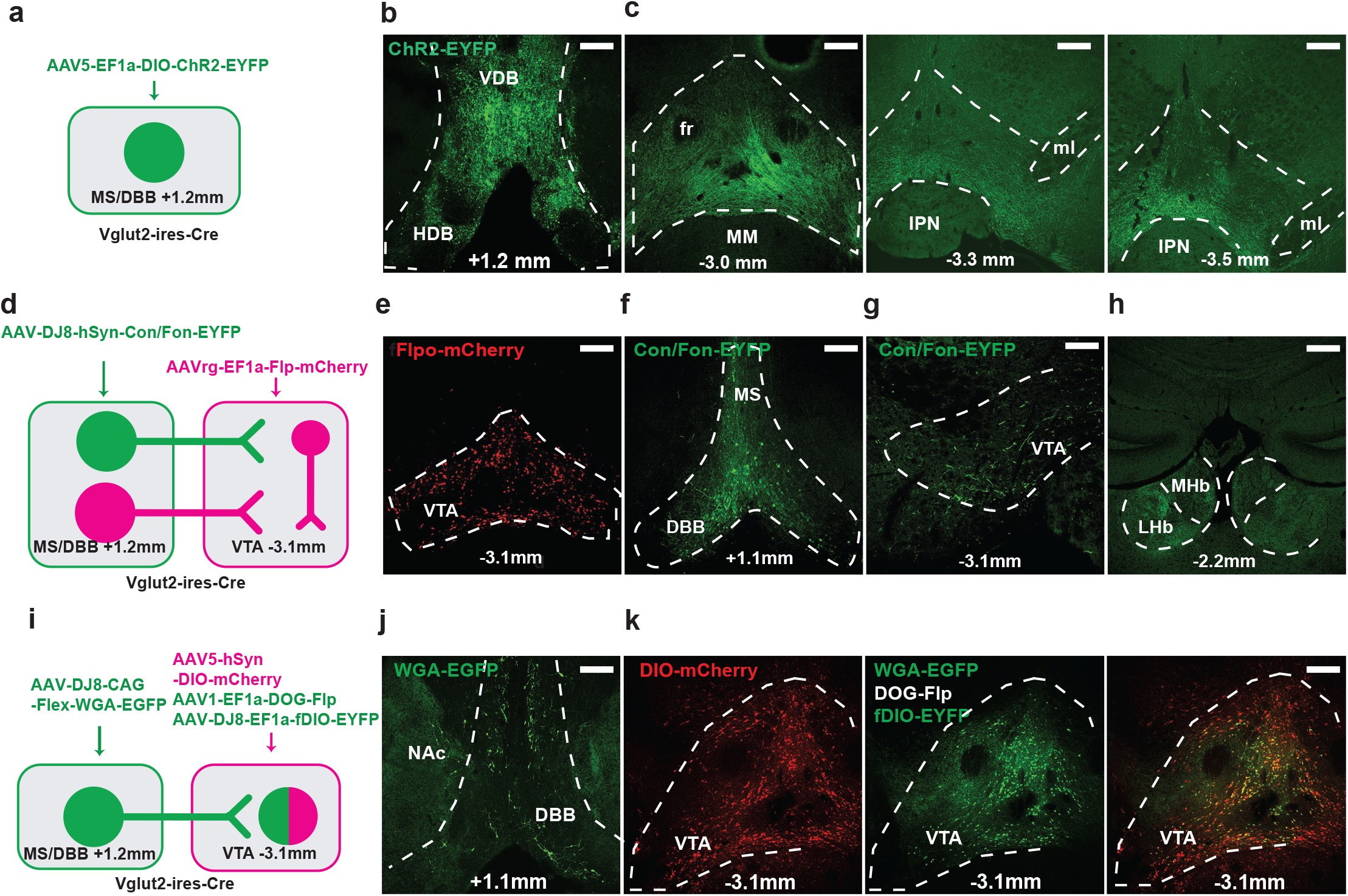
Direct projections from BF^glu+^ neurons to VTA^glu+^ neurons. a, Schematic diagram showing virus injection strategies for labeling BF downstream glu+ neuronal fibers. b, Expression pattern of the injected ChR2 virus in the BF. c, Expression pattern of ChR2-EYFP within the anterior to posterior sections of the VTA. d, Schematic diagram showing virus injections to label BF^glu+^ neurons that send projections to the VTA. e, Expression pattern of the Flp-mCherry virus in the VTA. f, Expression pattern of the injected Con/Fon-EGFP virus in the BF. EGFP labeled VTA-projecting BF^glu+^ neurons. g,h, EGFP-positive neuronal fibers from VTA-projecting BF^glu+^ neurons in the VTA (g) and the LHb (h). i, Schematic diagram showing virus injection strategies to label downstream VTA neurons that form synaptic connections with BF^glu+^ neurons. j, Expression pattern of the injected WGA-EGFP virus in the BF. k, Co-localization of EGFP positive neurons and mCherry neurons (glu+ neurons). Scale bar = 200 µm. VDB: vertical diagonal band of broca. HDB: horizontal diagonal band of broca. Fr: fasciculus retroflexus. MM: mammillary nucleus. IPN: interpeduncular nucleus. Ml: medial mammillary nucleus, lateral.

Given the heterogeneity of VTA neurons, we next sought to identify the neuronal subtype in the VTA that receives projections from BF^glu+^ neurons using the trans-synaptic tracer wheat germ agglutinin (WGA)^37^. For this, we delivered conditional AAV-DJ8-Flex-WGA-EGFP viral particles to the BF of Vglut2-ires-Cre mice (Figures 1i and 1j). EGFP signals that were traced from the BF were further amplified by co-delivery of AAV1-Ef1a-Flp-DOG-NW and AAV-DJ8-Ef1a-fDIO-EYFP in the VTA (Figures 1i and 1k). The AAV1-Ef1a-Flp-DOG-NW vector expresses Flp dependent on EGFP signals^38^, which in this case labeled VTA neurons that form synaptic contacts with BF^glu+^ neurons with EGFP expression. AAV5-hSyn-DIO-mCherry was delivered to the VTA to visualize glu+ neurons. As a result, abundant WGA-EGFP labeled neurons were co-localized with mCherry positive glu+ neurons as well as mCherry negative non-glu+ neurons (Figures 1k and S2). Since WGA is known to trace in both anterograde and retrograde fashions, we next used nontoxic tetanus toxin C-fragment (TTC)-based transsynaptic retrograde tracing to determine whether the VTA sends projections to BF^glu+^ neurons^39^. Toward this, we delivered AAV-DJ8-CAG-DIO-TTC-EGFP to the BF and a mixture of AAV5-hSyn-DIO-mCherry, AAV1-Ef1a-Flp-DOG-NW, and AAV-DJ8-Ef1a-fDIO-EYFP to the VTA of Vglut2-ires-Cre mice (Figure S3). Through this we identified EYFP expression in a subset of VTA neurons, and the vast majority of EYFP positive neurons were not mCherry positive (Figure S3b), suggesting VTA non-glu+ neurons, including DA and GABAERGIC neurons, send direct projections to the BF. In addition, VTA non-glu+ neurons displayed minimal co-localization of c-Fos expression upon photostimulation of the BF^glu+^ ➔ VTA circuit (Figure 4). These observations suggest that glu+ neurons in the VTA are the major downstream targets of BF^glu+^ neurons.

**Figure 2:**
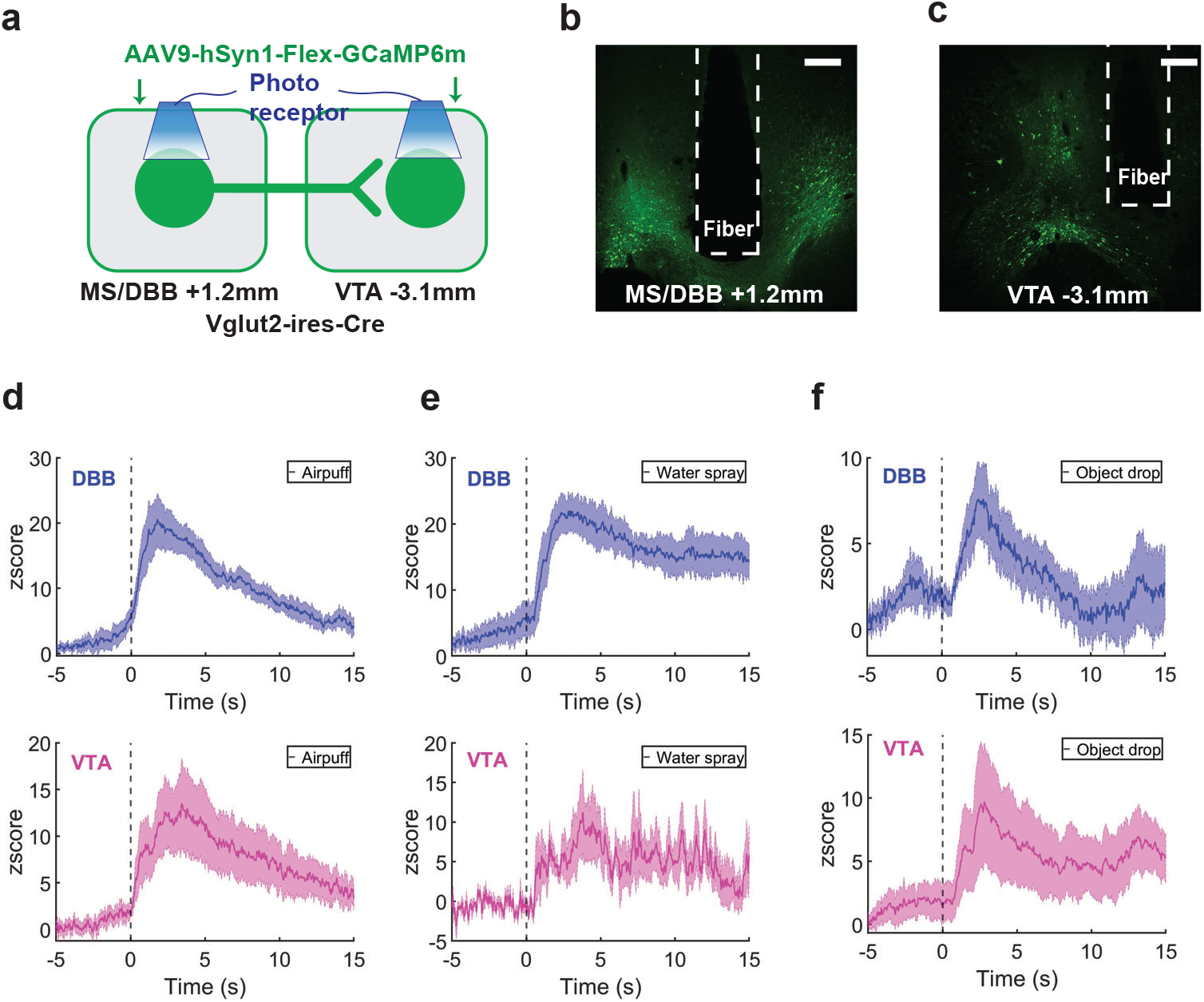
BF^glu+^ and VTA^glu+^ neurons showed correlated responses to external threat cues. a, Schematic diagram showing virus injections and optic cannula implantation in both the BF and the VTA for simultaneous dual GCaMP6m-based fiber photometry recordings. b,c, Expression patterns of the injected GCaMP6m virus and cannula tracks in the BF (b) and the VTA (c). Scale bar = 200 µm. d-f, BF (top panels) and VTA (bottom panels) Ca2+ signal Z scores in response to different physical stressors including air puff (d), water spray (e), and object drop (f). Time = 0 represents the starting time of the stimuli. The shades represent +/- SEM. Animals N = 7.

To examine whether VTA^glu+^ projecting BF neurons send collaterals to other brain sites, we next delivered AAV-DJ8-CAG-DIO-TTC-EGFP to the VTA and a mixture of AAV1-Ef1a-Flp-DOG-NW, AAV-DJ8-Ef1a-fDIO-EYFP and AAV8-Ef1a-Con/Fon-mCherry in the BF of Vglut2-ires-Cre mice (Figures S5a and S5b). In this configuration, EYFP expression will identify BF neurons that project to VTA^glu+^ neurons and mCherry expression will identify BF^glu+^ neurons that send direct projections to VTA^glu+^ neurons. Through this approach, we found both BF^glu+^ and BF non-glu+ neurons were traced (Figure S5b), suggesting both groups send direct projections to VTA^glu+^ neurons. In addition, Con/Fon-mCherry positive fibers from VTA^glu+^ neurons projecting BF^glu+^ neurons were also detected in other brain regions (Figures S5c and S5d), revealing patterns of collateralization.

### Correlated responses of BF^glu+^ and VTA^glu+^ neurons to environmental cues

BF^glu+^ neurons respond to various odor stimuli, including both aversive and food odors^15^. However, how these neurons transmit these signals to other brain regions is unknown. VTA^glu+^ neurons have also been shown to be sensitive to aversive stimuli^40–42^. To study the functional relationship between BF^glu+^ and VTA^glu+^ neurons in responding to environmental stressors, we performed GCaMP6m-based *in vivo* dual-fiber photometry recordings in both groups of neurons in freely moving mice. We delivered AAV9-CAG-Flex-GCaMP6m viral particles to both the BF and the VTA of Vglut2-ires-Cre mice, and implanted optic cannulas independently targeting both areas (Figure 2a). The expression of GCaMP6m was confirmed in glu+ neurons of the BF and VTA (Figures 2b and 2c). As expected, BF^glu+^ neurons exhibited an increased activity to air puff, water spray, and object drop (Figures 2c-2f). VTA^glu+^ neurons also exhibited reliable and correlative activities similar to BF^glu+^ neurons (Figures 2d-2f), which is consistent with a direct excitatory glutamatergic projection from the BF to VTA^glu+^ neurons.

### Activation of BF^glu+^ **➔**VTA circuit reduces feeding and causes avoidance

Heightened activity of BF^glu+^ neurons is known to inhibit food intake^15^. Given that we revealed BF^glu+^ ➔ VTA^glu+^ connectivity, we reasoned that VTA^glu+^ neurons may mediate the effects on feeding. To test this, we utilized ChR2-based *in vivo* optogenetics to activate BF^glu+^ neuronal fibers within the VTA (Figure S3a). Toward this, we delivered AAV5-Ef1a-Flex-ChR2-EYFP in the BF of Vglut2-ires-Cre mice and implanted fiber optic cannulas over the VTA. Photostimulation (λ = 473 nm, 20 HZ-20 ms, 10 mins) of ChR2 positive fibers within the VTA was found to significantly increase c-Fos in a subset of VTA neurons (Figures 3b and 3c), confirming functional activation of these neurons. Next, in fasted mice, *in vivo* photostimulation of the BF^glu+^ ➔ VTA circuit reduced food consumption (Figure 3d). This effect was partially rescued by the administration of glutamate receptor antagonists AP5 and DNQX into the VTA (Figure 3d), suggesting that the reduction in feeding was mediated by activation of glutamate receptors in the VTA. To test the valence associated with the observed feeding inhibition, a real-time place avoidance test (RTPA) was conducted, in which the photostimulation was paired with one half of the arena in a counterbalanced manner. Mice expressing ChR2 spent more time in the non-stimulation half, while EYFP-expressing controls showed no preference for either side, suggesting an avoidance behavior elicited by activation of the BF^glu+^➔VTA circuit (Figures 3e and 3f). Interestingly, ChR2-expressing mice also exhibited an increased physical activity level with photostimulation (Figure 3f), which was corroborated by increased locomotion during the open field test (Figures 3g and 3h). Since VTA-projecting BF^glu+^ neurons also send collateral projections to the LHb and other brain regions, the observed behavioral effects may be partly mediated by these additional projection sites through parallel activations. To test this possibility, we performed the same experiment but with additional simultaneous inhibition of BF^glu+^ neuron soma, which would largely eliminate collateral activation. We expressed the inhibitory designer receptor exclusively activated by designer drug (Gi, hM4Di) in BF^glu+^ neurons (Figure 3i). CNO treatment effectively reduced c-Fos expression in BF^glu+^ neurons induced by activation of glu+ fibers in the VTA (Figure 3j), confirming an effective inhibition of BF^glu+^ neurons. Behaviorally, with the presence of CNO, photostimulation of BF^glu+^ fibers in the VTA was still able to induce aversion, increase physical activity and reduce fasting refeeding (Figure 3k and 3i), suggesting that the observed effects were mediated by BF^glu+^ projections within the VTA.

**Figure 3.**
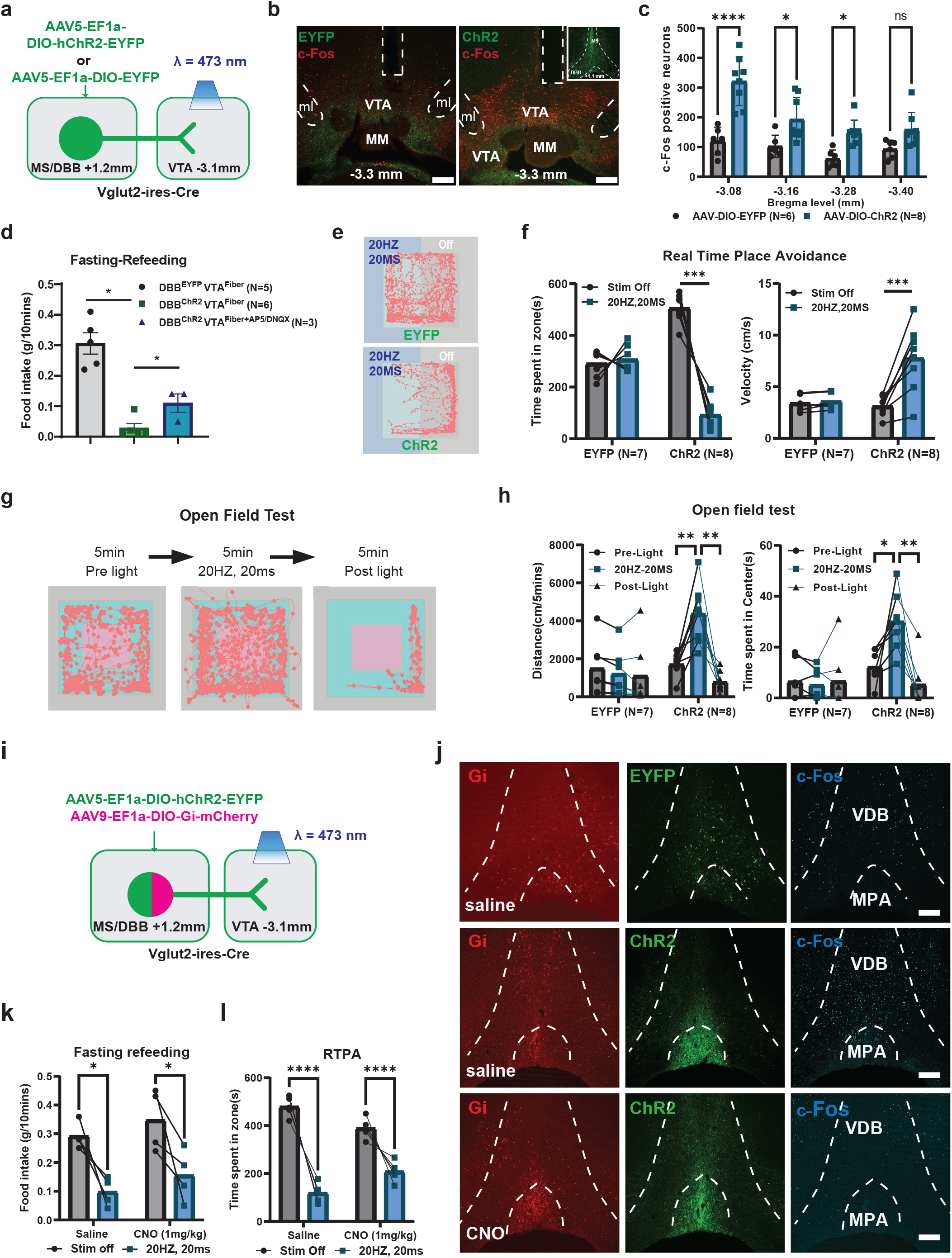
*In vivo* activation of BF^glu+^ ➔ VTA^glu+^ projections suppressed food intake. a. Schematic diagram showing strategies of virus injections and optic cannula implantations for photostimulation of BF^glu+^ ➔ VTA^glu+^ projections. b, Representative expression patterns of the injected ChR2 virus and cannula tracks in the VTA. C-Fos is a marker of neuronal activation. Scale bar = 200 µm. c, Quantitative comparisons of c-Fos positive neuron numbers at different bregma levels (from anterior to posterior) between control EYFP and ChR2 groups. Two-way ANOVA followed by Sidak multiple comparisons test; from anterior to posterior bregma levels: F (1, 48) = 56.18; P < 0.0001. d, The comparison in food intake induced by overnight fasting between the two groups. Unpaired student’s t-tests: ChR2 vs, EYFP, P <0.0001; ChR2+AP5/DNQX vs. ChR2, P = 0.0350. e, Representative moving tracks (pink color) of animals from EYFP (top) and ChR2 (bottom) group in the real time place avoidance test. The left half of chamber (blue and light cyan color) was paired with photostimulation (20 HZ, 20 MS) while the right half (gray color) was not. f, Quantitative comparisons in duration and velocity in two halves of chambers between control EYFP and ChR2 groups. Two-way ANOVA followed by Sidak multiple comparisons test: Duration, F (1, 13) = 61.16, P < 0.0001; Velocity, F (1, 12) = 16.83, P = 0.0015. g, 5-min moving tracks (pink traces) of a representative animal from the ChR2 group in pre-light, light-on and post-light conditions. Pink: center area. Cyan: peripheral area. Gray: wall. h, Quantitative comparisons in distance and time in the center in the open field test (OFT) between the two groups. Two-way ANOVA followed by Sidak multiple comparisons test: Distance, F (2, 26) = 20.71, P < 0.0001; Time in center, F (2, 24) = 11.60, P = 0.0003. i, Schematic diagram showing virus injections and optic cannula implantation to test the behavioral effects of inhibiting co-lateral projections from VTA projecting BF^glu+^ neurons. j, c-Fos expression patterns in the BF. CNO application reduced the c-Fos activation induced by the photostimulation. k, Quantification results of fasting refeeding test and real time place avoidance test. Two-way ANOVA followed by Sidak multiple comparisons test: Fasting refeeding, F (1, 6) = 0.0008487, P = 0.9777; RTPA, F (1, 8) = 23.61, P = 0.0013.

### Activation of VTA^glu+^ neurons reduces motivational feeding and body weight, and increases activity

Given that VTA^glu+^ neurons are one of major downstream targets of BF^glu+^ neurons, we next examined the role of VTA^glu+^ neurons in feeding regulation. Toward this, we first employed a Gq (hM3Dq)-dependent chemogenetic method to activate VTA^glu+^ neurons in a short term and assessed the effects (Figure 4a). We delivered AAV9-hSyn-DIO-Gq-mCherry or AAV5-hSyn-DIO-mCherry viral particles to the VTA of Vglut2-ires-cre mice. Notably, the administration of the Gq agonist CNO (i.p., 1mg/kg) significantly increased c-Fos expression in Gq-mCherry-expressing neurons comparing to the mCherry group (Figure 4b and 4c). Also, CNO administration in the Gq group greatly reduced fasting-induced refeeding compared to saline, while having no effects in the mCherry group within 6 hours after treatments (Figure 4d). Given the role of VTA^glu+^ neurons in reward processing^43,44^, we further explored the effects of these neurons in food motivation. For this, mice were trained through an established protocol to perform nose pokes at the correct port for sucrose pellets with increasing fixed ratios (FRs) until they reached the point of acquiring more than 20 sucrose pellets at FR = 15 within a 30-minute session (Figure 4e). During the testing episode, CNO administration completely blocked nose poke behaviors for sucrose pellets in the Gq group, while having no effects on the mCherry control group (Figure 4f). These results demonstrated a profound effect of VTA^glu+^ neuron activation on suppressing motivational feeding behaviors. Consistent with the increase in physical activity by *in vivo* photostimulation of the BF^glu+^ ➔ VTA circuit, CNO-mediated activation of VTA^glu+^ neurons increased physical activity measured in metabolic cages (Figures 4g and 4h). Since changes in feeding are known to be associated with altered anxiety ^1^, we further assessed the anxiety level in these mice using the light-dark box test (LDT). Compared to saline treatments, CNO application in the Gq group reduced time spent in the light side of the box, while CNO had no effects in the mCherry group (Figure 4i), suggesting that CNO-induced activation of VTA^glu+^ neurons induced anxiety-like behavior. In sum, activation of VTA^glu+^ neurons led to increased locomotion, increased anxiety-like behaviors, and hypophagia with reduced motivation for food, all of which are reminiscent of typical phenotypes in anorexia patients, suggesting a role of these neurons in the pathogenesis of anorexia.

**Figure 4.**
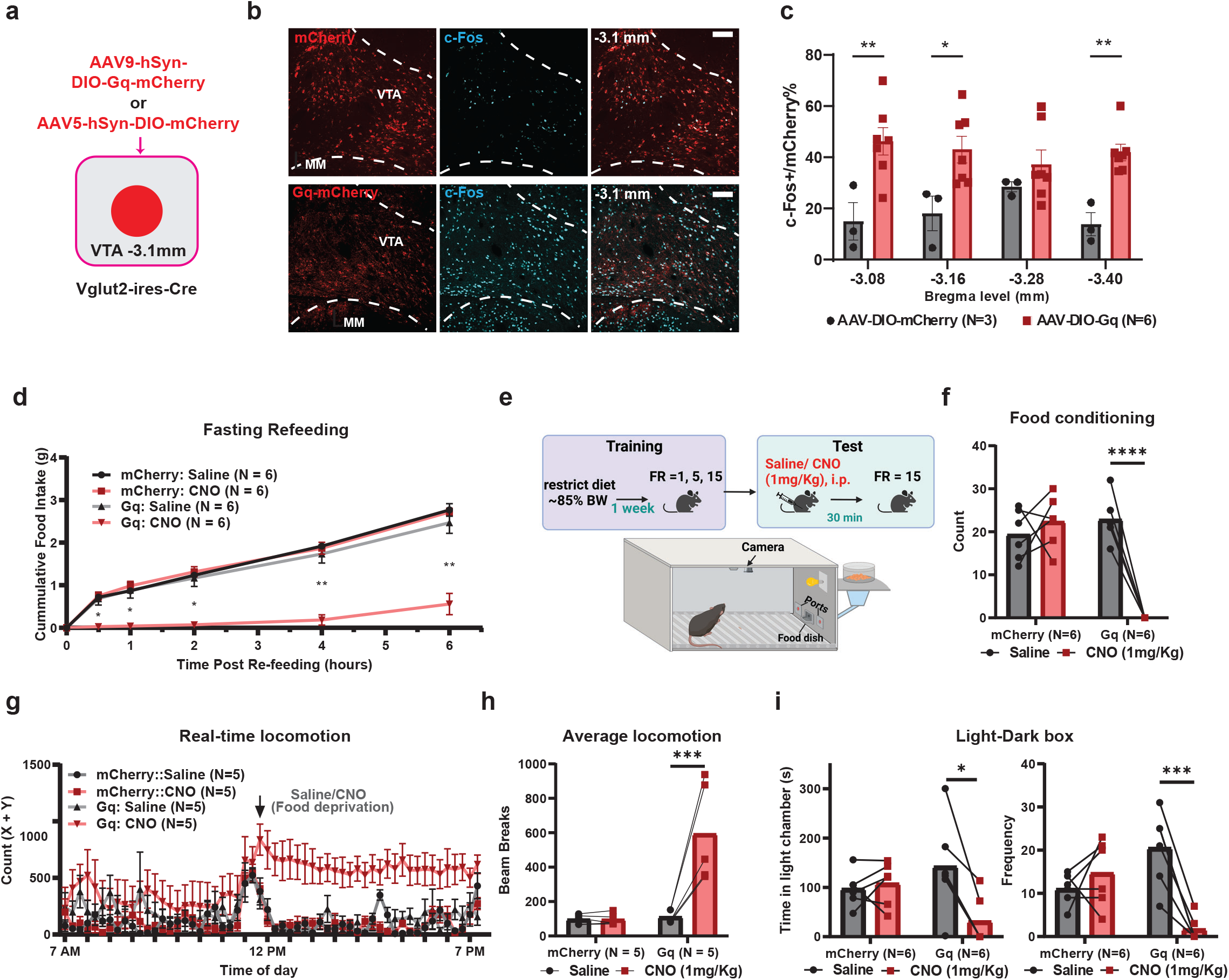
Acute activation of VTA^glu+^ neurons decreased food motivation and induces anxiety. a, Schematic diagram showing the Gq virus injection in VTA^glu+^ neurons for chemogenetic activation. b, Expression pattern of mCherry (top) or Gq (bottom) and c-Fos immunostaining in the VTA. Scale bar = 200 µm. c, Qualitative comparisons in the percentages of c-Fos positive neurons in the viral (mCherry) labelled neurons shown in an anterior to posterior bregma order. Two-way repeated ANOVA followed by Sidak multiple comparisons test: F (1,32) = 30.24, P < 0.0001. d, Comparisons in cumulative fasting-refeeding food intake within 6 hours post saline and CNO application between mCherry and Gq groups. Two-way ANOVA followed by Sidak multiple comparisons test: F (15, 105) = 19.90, P < 0.0001. e, Schematic diagram showing procedures and timing of nose poke training and testing in mice. FR: fixed ratio. f, Comparisons in sucrose pellets acquired during the testing session between the mCherry and Gq groups after saline and CNO application. Two-way ANOVA followed by Sidak multiple comparisons test: F (1, 10) = 56.42, P < 0.0001. g, Real-time locomotor activities from metabolic cages. The arrow indicates when the mice were injected with saline or CNO. h, Comparisons of average locomotion within 6 hours after saline or CNO treatment. Two-way repeated ANOVA followed by Sidak multiple comparisons test: F (1, 8) = 17.76, P = 0.0029. i. Comparisons of time (left) spent and frequency entering (right) in light chambers during the light-dark box test. Two-way repeated ANOVA followed by Sidak multiple comparisons test: Time, F (1, 10) = 11.41, P = 0.007; Frequency, F (1, 10) = 21.39, P = 0.0009.

Since anorexia symptoms are likely caused by chronic changes in neuronal activity, we next explored the effect of chronic activation of VTA^glu+^ neurons utilizing a mutated sodium channel from bacteria (NaChBac) that enables long-term activation^45,46^. We delivered to the VTA of Vglut2-ires-Cre mice with AAV-DJ8-Ef1a-DIO-NaChBac-EGFP or control AAV5-Ef1a-DIO-EYFP viral vectors (Figure 5a). Immunostaining results showed that NaChBac expression led to a dramatic increase in c-Fos expression in the VTA, confirming the activation of VTA^glu+^ neurons (Figures 5b and 5c). Interestingly, both male and female mice with NaChBac expression showed lower body weight and exhibited resistance to diet-induced obesity, compared to the EYFP-injected control group (Figures 5d and S7). Consistent with lower body weight, NaChBac group displayed lower food intake compared to controls (Figure 5e). When measured in metabolic cages to collect the real-time physiological data, compared to controls, NaChBac group showed increased locomotion (Figures 5f and 5g), suggesting that the lower body weight was due to further energy deficits. Together, these results demonstrate that chronic activation of VTA^glu+^ neurons causes hypophagia, decreased body weight gain and higher locomotion, consistent with typical symptoms observed in anorexia patients.

**Figure 5.**
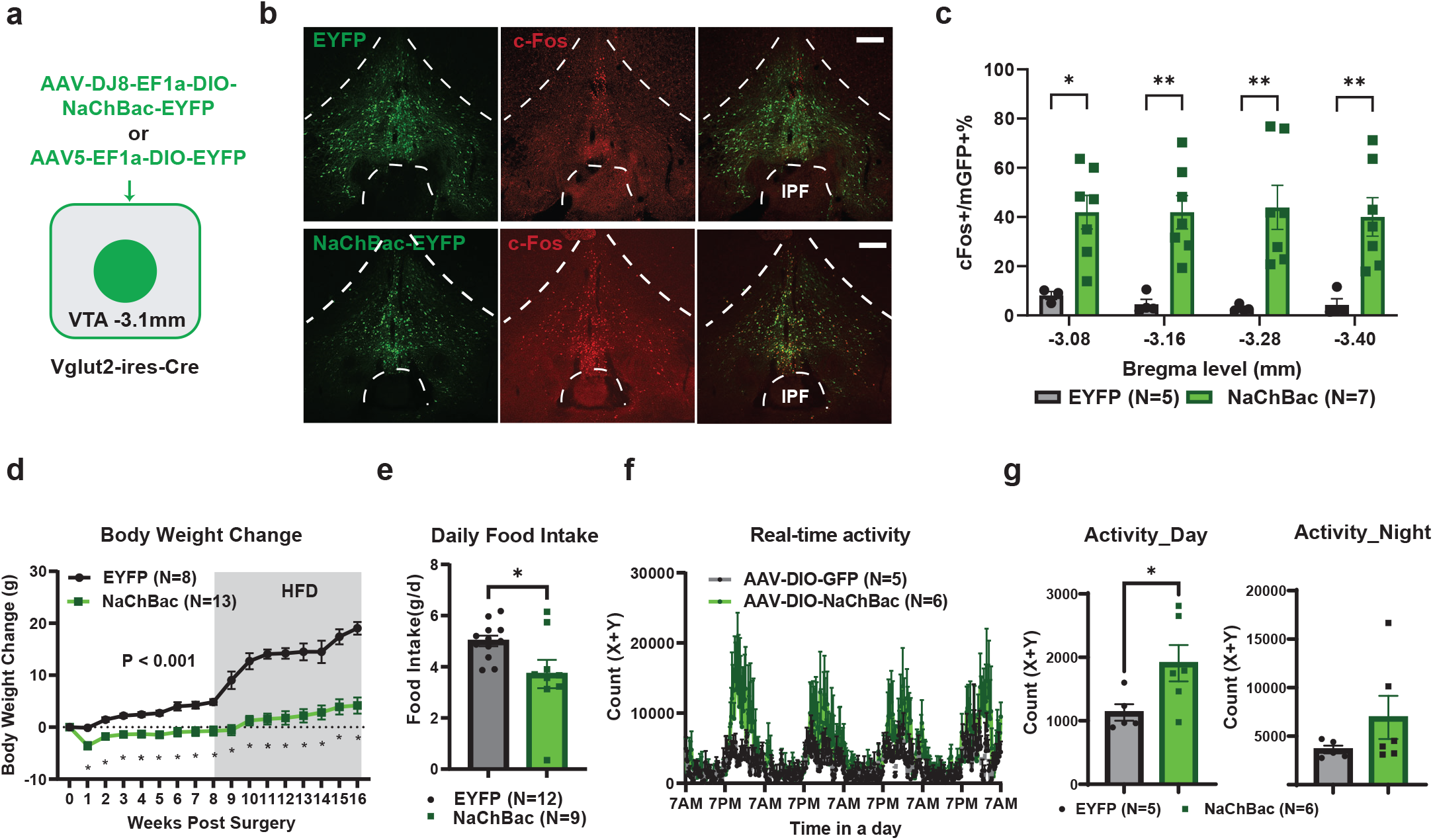
Effects of chronic activation of VTA^glu+^ neurons led to major hallmarks of anorexia. a, Schematic diagram showing injections of the NaChBac virus to the VTA for chronic activation of VTA^glu+^ neurons. b, Representative coronal sections of the VTA showing c-Fos immunostaining from EYFP (top) and NaChBac (bottom) mice. Scale bar = 200 µm. c, Quantitative comparisons in the number of c-Fos positive neurons in EGFP-labeled glu+ neurons between control and NachBac mice. Two-way ANOVA followed by Sidak multiple comparisons test: F (1, 35) = 46.51, P < 0.0001. d, Comparisons in weekly body weight for the 16 weeks after viral delivery. From 0 to 8 weeks post-surgery, mice were fed chow diet and from 9 to 16 weeks post-surgery, mice were fed with high fat diet (HFD). Two-way repeated ANOVA followed by Sidak multiple comparisons test: F (16, 190) = 17.22, P < 0.0001. e, The comparison in daily food intake on chow diet between the two groups. Unpaired student’s t test, P = 0.0251. f, Real-time locomotor activity patterns measured by the CLAMS metabolic cages. g, Comparisons in the locomotor activity levels measured during periods of the day (left) and the night (right). Unpaired t-test: Day time, P = 0.0475; Night time, P = 0.2154.

### The BF^glu+^ **➔** VTA^glu+^ projection suppresses DA release

Changes in DA signaling have been associated with lower body weight^47^. Given the newly-revealed role of the BF^glu+^ ➔ VTA^glu+^ circuit in motivational feeding, we further investigated potential effects of this projection in modulating DA release. The NAc is the major dopaminergic reward center, and one of the major downstream regions for VT DA neurons, so we recorded dynamic DA release in the NAc using gGRAB-DA3m (gDA3m)-based fiber photometry recording in freely moving mice (manuscript submitted). For this, we delivered an AAV9-hSyn-gDA3m virus to the shell of NAc (NAcSh) and implanted optic fibers cannulas targeting the same region (Figures 6a and 6b). Consistent with previous results in sucrose consumption^29,48^, we detected an increase in DA release in the NAcSh when mice consumed a high-fat diet pellet (Figures 6c-6e). In addition, we observed reduced DA release when mice received an aversive stimulus such as water spray (Figure 6f-6h). Interestingly, we observed a rebound in DA release following the suppression period induced by aversive stimuli (Figures 6f-6h), reminiscent of the comforting effects of the “pain relief” observed previously^49^.

**Figure 6.**
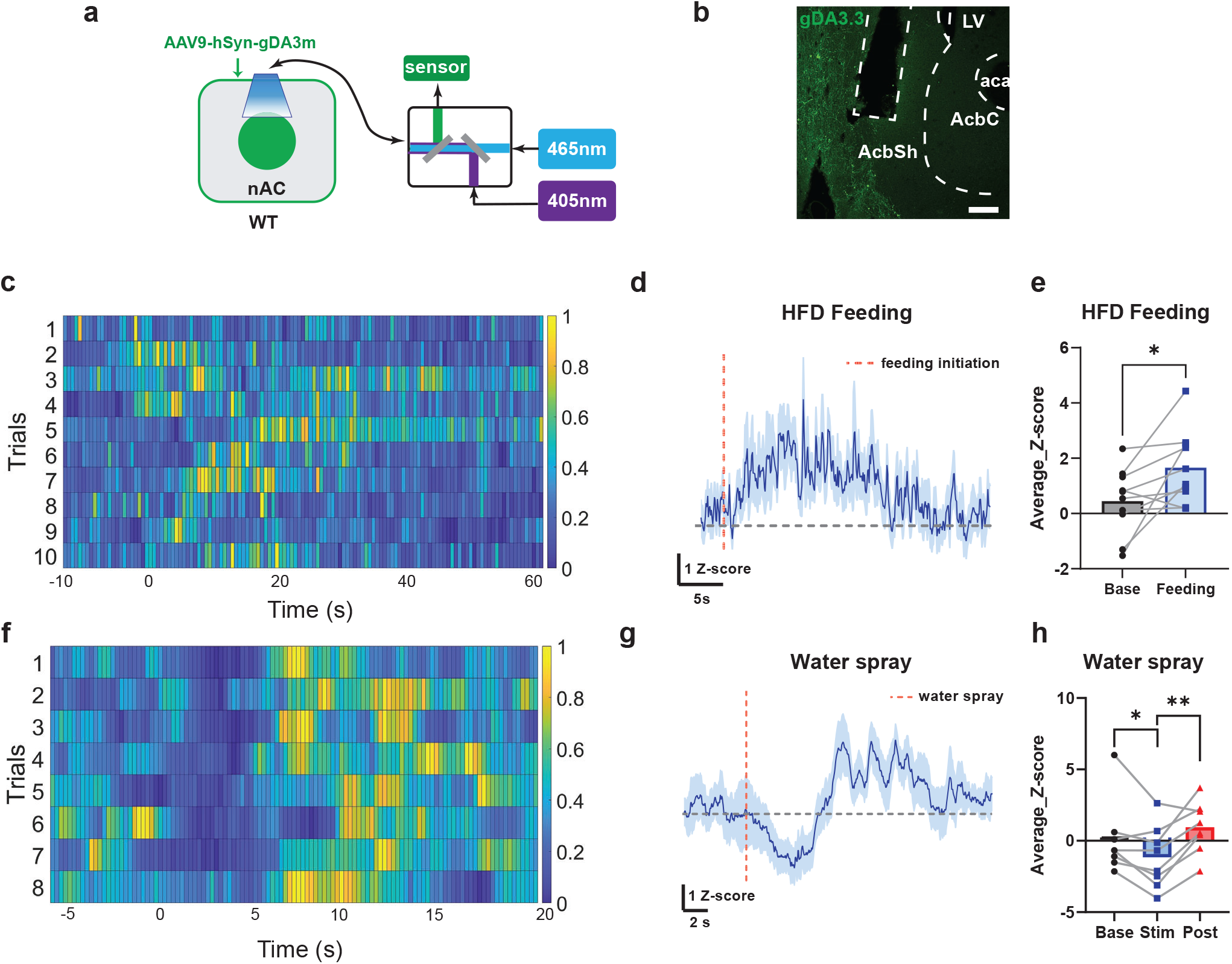
Changes of DA release in the NAc during feeding and physical stresses. a, Schematic diagram showing virus injections and optic cannula implantations for recording DA release in the NAc. b, Representative coronal sections of the VTA showing gDA3m expression pattern and cannula tracks. Scale bar = 200 µm. c, Heatmap of DA release Z-score signals corresponding to first feeding bouts of individual trials. Signals were re-scaled from 0 to 1 across each row. d, Averaged Z-score signals of DA release in the NAc corresponding to first feeding bouts. The light blue shade represents signals with mean +/- SEM. Time = 0 was the onset of feeding. e, Averaged Z-scores during baseline and feeding. Paired student’s t-test: P = 0.0268. f, Heatmap of signals of DA release Z-score signals of individual trials in response to water spray. Signals were re-scaled from 0 to 1 across each row. g, Averaged signals of DA release Z-score signals in the NAc in response to water spray. The light blue shade represents signals with +/- SEM. Time = 0 indicating the onset of water spray to mice. h, Averaged Z-scores during baseline, 5 seconds post water spray, and 5 to 10 seconds post water spray. Paired student t-test: Stim vs. Base, P = 0.0319; Post vs. Stim, P = 0.0032. Base: baseline. Feeding: feeding bout. Stim: 5 seconds post water spray. Post: 5 to 10 seconds post water spray.

Finally, to examine the effect of BF^glu+^ ➔ VTA^glu+^ projections on DA release, we recorded DA release while activating BF^glu+^ fibers in the VTA via targeted photostimulation (Figure 7a). Toward this, we delivered AAV5-Ef1a-DIO-ChR2-EYFP in the BF of Vglut2-ires-Cre mice and implanted optic cannulas in the VTA in addition to delivering gDA3m virus and implanted optic fibers in the NAcSh (Figures 7a-7d). Photostimulation of the BF➔VTA circuit with laser at 20HZ-20ms for durations of 10s and 20s both reduced DA release in a duration-dependent manner, which was again followed with a rebound DA phase of release (Figures 7e-7g, and S8). To explore the relationship between DA release regulated by the BF^glu+^ ➔ VTA^glu+^ projection and feeding behaviors, we monitored DA release in response to photostimulation when mice initiated HFD feeding bouts. In this regard, *ad libitum* mice were allowed to move freely to reach a HFD pellet placed in one corner of the cage. At the time point when the mice started consuming pellets, photostimulation was applied to activate the BF^glu+^ VTA^glu+^ projection for a duration of 10 seconds (Figure 7i). The signals for DA release increased when mice were involved in pellet consumption, and were reduced upon photostimulation, which was also associated with an immediate pause on feeding and a roam away from the food (Figures 7h-7j). These results collectively suggest that activation of BF^glu+^ ➔ VTA^glu+^ projection reduces feeding and involves reduced DA release in the NAc.

**Figure 7.**
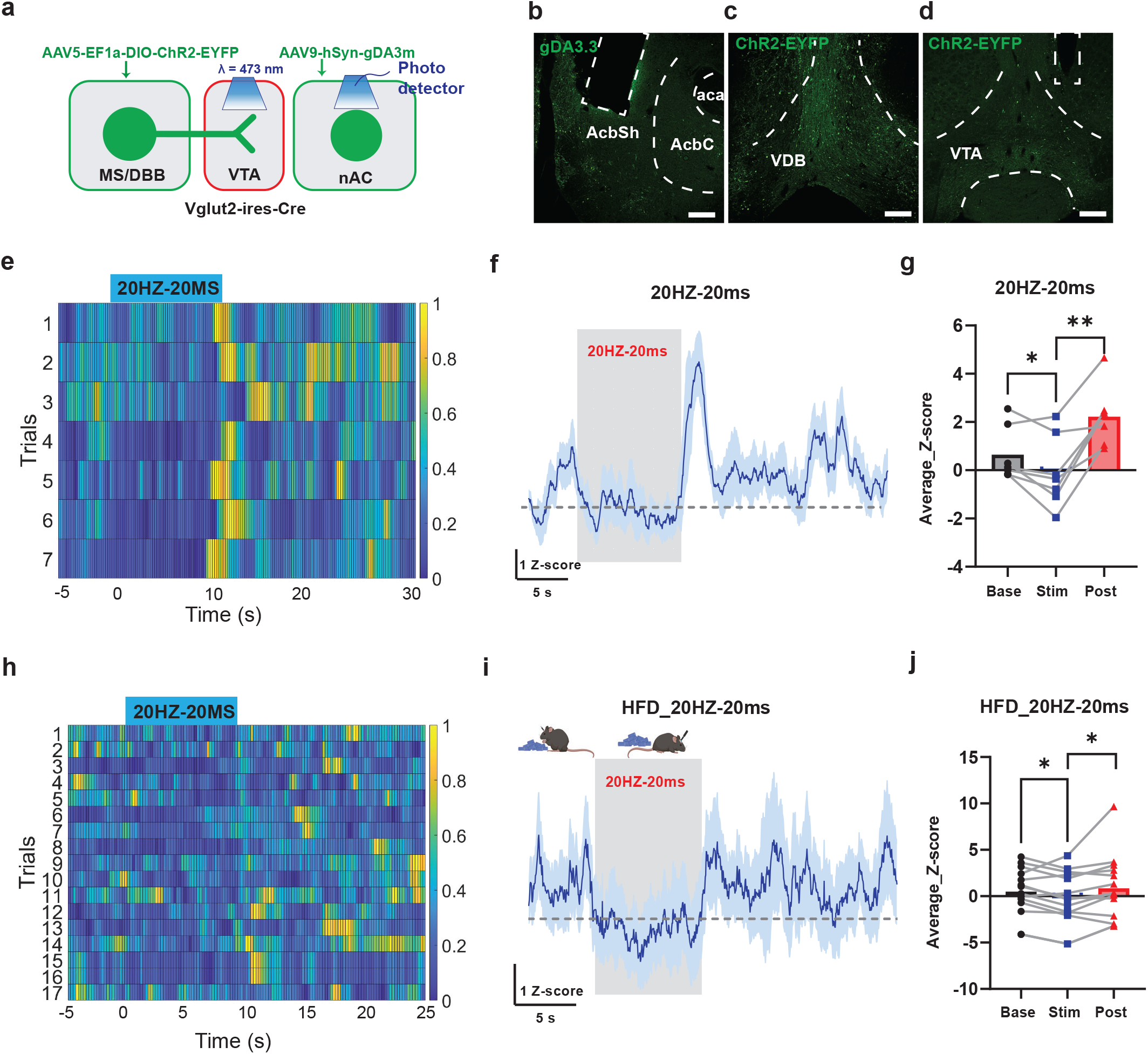
Changes in DA release in the NAc in response to activation of the BF^glu+^ ➔ VTA^glu+^ projections. a, Schematic diagram showing virus injections and optic cannula implantations to record DA release in the NAc with photostimulation of BF^glu+^ ➔ VTA^glu+^ projections. b-d, Representative coronal sections of the BF, the VTA and the NAc showing injection patterns of ChR2 and gDA3m and cannula tracks. Scale bar = 200 µm. e, Heatmap of DA release Z-score signals of individual trials in response to photostimulation with 10-second duration. Signals were re-scaled from 0 to 1 across each row. The indigo box indicates the period of photostimulation. f, Averaged trace of DA release Z-score signals in the NAc in response to a period of 10 seconds of 20HZ-20ms photostimulation. g, Averaged Z-scores during baseline, 20HZ-20ms stimulation and 5 seconds post stimulation. Paired student t-test: Stim vs. Base, P = 0.0349; Post vs. Stim, P = 0.0019. h, Heatmap of DA release Z-scores signals of individual trials when mice were engaging HFD feeding with photostimulation. Signals were re-scaled from 0 to 1 across each row. The indigo box shows the period with photostimulation. i, Averaged signals of DA release in the NAc in response to a period of 10 seconds of 20HZ-20ms photostimulation right after the onset of HFD feeding. The blue shade represents signals in +/- SEM. Time = 0 was the onset of photostimulation. j, Averaged Z-scores during baseline, 20HZ-20ms photostimulation, and 5 seconds post stimulation. Paired student t-test: Stim vs. Base, P = 0.0213; Post vs. Stim, P = 0.0223. Base: baseline. Stim: 20HZ-20ms photostimulation. Post: 5 seconds post photostimulation.

## Discussion

An appropriate decision to engage in feeding is an adaptive behavior that integrates perceived stress and internal energy needs. Environmental disturbances cause stress and limit feeding, even when mice have great energy demands. Several studies on stress-induced hypophagia have focused on neural pathways that process sensory cues for threats, which also elicit behavioral signs of stress and inhibit feeding ^1,2^. In this study, we have identified a BF^glu+^ ➔ VTA^glu+^ circuit that is sensitive to and activated by environmental cues, which is associated with reduced DA release in the NAc. *In vivo* activation of this circuit reduced feeding, increased physical activity, and caused avoidance behaviors. Similarly, activation of downstream VTA^glu+^ neurons diminished food motivation, caused hyperactivity, and increased behavioral signs of anxiety. Consistently, mice with chronic activation of VTA^glu+^ neurons exhibited hypophagia and reduced body weight associated with heightened physical activity. Interestingly, the observed phenotypes represent typical behavioral and physiological signs observed in patients with anorexia, the core symptoms of which include voluntary starvation, hyperactivity, and anxiety^5,6,50^. Taken together, our results reveal that overactivation of the BF^glu+^ ➔ VTA^glu+^ circuit may contribute to the pathogenesis of anorexia.

Consistent with previous observations that BF neurons responds to volatile odors and predator cues^15,17,18^, here we showed that BF^glu+^ neurons were sensitive to various physical stressors and aversive stimuli. To extend these findings, we also found that VTA^glu+^ neurons are directly downstream targets of BF^glu+^ neurons, and that these VTA neurons also respond to aversive stimuli in correlation, suggesting that they are important downstream mediators in aversive sensory perception. Our previous results showed that chronic activation of BF^glu+^ neurons elicited a starvation phenotype^15^, which supports our current observations that activation of the BF^glu+^ ➔ VTA^glu+^ circuitry causes an anorexia-like behaviors. Taken together, these observations support that BF^glu+^ neurons function as a primary node in the brain that senses external threat cues and overrides energy needs to avoid potential risks. Since the VTA is known to be involved in motivation and feeding, the BF^glu+^➔ VTA^glu+^ circuit is well positioned to integrate sensory cues and impact motivational feeding behaviors. It is worth noting that the VTA-projecting BF^glu+^ neurons also send collaterals to other brain regions including lateral habenula (LHb) and lateral hypothalamus (LH) neurons. Previous studies suggest that both LH and LHb neurons mediate the action of BF^glu+^ neurons in feeding inhibition^15,17^. Since both the LH and LHb have been shown to project to the VTA^51,52^, the VTA represents a convergent brain target from various brain sites in mediating adaptive feeding behaviors to ongoing environmental threat cues.

Despite extensive studies on the pathogenesis of anorexia, the underlying brain mechanism remains elusive. One of the major difficulties in this line of research lies in the lack of reliable animal models that could closely capture the symptoms observed in human anorexia patients^53^. Regarding this, chronic activation of BF^glu+^ neurons has been shown to cause a starvation phenotype^15^. Consistently, our current results demonstrate that chronic activation of VTA^glu+^ neurons, as one of the downstream targets of BF^glu+^ neurons, led to reduced motivation for feeding, heightened locomotion, and diminished HFD-induced obesity. These observations, taken together, support that uncontrolled overactivation of the BF^glu+^ ➔ VTA^glu+^ circuit may contribute to anorexia-like phenotypes. Given the role of BF^glu+^ neurons in sensing environmental cues^15^, it is conceivable that hypersensitivity by these neurons to environmental threats may cause overactivation. Given their function in memory and conditioning^7,54^, activation of BF neurons may also be caused by various conditioned cues, which may also lead to a predisposition to the development of anorexia. In particular, chemogenetic activation of VTA^glu+^ neurons caused anxiety-like behaviors, which is in line with previous studies showing that sustained stimulation of VTA^glu+^ neurons is less preferred and causes behavioral avoidance^32,55^. This effect can be mediated through projections to the LHb or to GABAergic neurons within the NAc^32,55^. Alternatively, since a subset of VTA^glu+^ neurons have been shown to co-release GABA^56^, GABA action from these neurons may also contribute to the observed effect since VTA GABAergic neurons can cause aversive responses ^30^.

Our results also showed that activation of the BF^glu+^ ➔ VTA^glu+^ circuit in mice resulted in anxiety-like behaviors, which reduced appetite even in the presence of strong internal energy needs and was associated with reduced DA release in the NAc. Although it is unknown how activation of VTA^glu+^ neurons led to reduced DA release, recent studies suggest that VTA non-DA neurons, including glu+ neurons, can gate DA release and modulate reward-related behaviors^25,27^. One possibility is through long-range projections. For example, VTA^glu+^ neurons send projections to the LHb glu+ neurons^32^, which send further excitatory inputs to rostromedial tegmental nucleus GABA neurons^57–59^, and further inhibit DA release through inhibitory synapses with VTA DA neurons^60^. It is also possible that VTA^glu+^ neurons reduce DA release through projections to the striatum via local circuits^61^. Further studies need to be conducted to test the possibilities.

Despite that the discussed literature on VTA^glu+^ neurons in causing aversion is consistent with our findings, other studies suggest that direct optical activation of VTA^glu+^ neurons can induce appetitive operant conditioning and reinforcement in the absence of DA release^31,34,56^. Our finding that activation of the BF^glu+^ ➔ VTA^glu+^ circuit induced hyperactivity was associated with reduced DA release is not in line with the general belief that overactivation of dopamine pathways cause hyperactivity, or that loss of DA release causes hypoactivity^62–64^. However in these cases, the role of DA action on locomotion is thought to be from substantial nigra neurons instead of the VTA. Previous studies also suggest a role of VTA^glu+^ neurons in driving arousal, exploration and facilitating defensive escape behaviors, which are all associated with hyperactivity, supporting a role for these neurons in increasing locomotion^40,65^. Supporting this, glutamatergic action in the NAc has been shown to increase locomotion^66,67^. Given the complexity of VTA local and remote circuits, further studies are required to address these observed inconsistencies.

It is not clear whether and to which degree the observed reduction in DA release contributes to the hypophagia phenotype. It was previously shown that modulation of dopamine receptor D2 (DRD2) signaling in the NAc caused anorexia-like behaviors in mice^68^. In addition, changes of dopamine signaling were observed in patients diagnosed with anorexia^47,69^. Human studies suggest a negative correlation between levels of DRD2 levels and BMI^70^. Animal studies suggest that an increase in DA signals is generally associated with positive valence and reward or reward-related cues^29,48,71,72^, whereas complete loss of DA release leads to a lethal phenotype, which can be rescued by the recovery of feeding through increasing DA release in the striatum^73^. These observations support the possibility that the hypophagia caused by overactivation of the BF^glu+^ ➔ VTA^glu+^ circuit is mediated by reduction in DA release in the striatum.

In summary, our results presented here support the conclusion that the BF^glu+^ ➔ VTA^glu+^ circuit functions to sense environmental threats and adjusts adaptive behaviors in feeding during hunger-threat conflict situations. Overactivation of this circuit led to hypophagia, reduced motivation for feeding, hyperactivity, and anxiety-like behaviors, all typical signs of anorexia. In addition to glu+ neurons, the BF also contains GABA and cholinergic neurons, both of which are also sensitive to environmental cues for threats^9–11^, suggesting a general role for the BF in surveying external cues and mediating necessary adaptation for survival. Since BF^glu+^ neurons send projections to several brain sites that also reduce feeding, VTA^glu+^ neurons may represent one downstream neuronal population that mediate the adaptive behaviors elicited by cue-activated BF^glu+^ neurons^15,17^. Since VTA^glu+^ neurons have also been shown to receive multiple upstream inputs implicated in mediating behaviors that include sleep, defense and exercise^44,74^, in addition to feeding behaviors shown here, VTA^glu+^ neurons appear to serve as a hub integrating both external and internal cues to mediate adaptive feeding behaviors toward the ever-changing environment.

## Methods and material

### Animal care

Animal care and procedures were approved by the University of Texas Health Science Center Houston Institutional Animal Care and Use Committee. Mice were housed at 21–22□°C on a 12□h light/12□h dark cycle with standard pellet chow and water ad libitum otherwise noted for fasting experiments, calorie-restricted diet for nosepoke training or high fat diet treatment. Vglut2-ires-Cre and Vgat-ires-Flp mice were purchased from The Jackson Laboratory (strain no. 016963 and no. 031331) and described previously^36,75^. Male and female mice were used in preliminary study and no significant difference was revealed. Mice used in experiments were acquired from same litters in different treatment groups and were 7∼16 weeks when used for surgery purposes.

### Steretotaxic surgery and viral vectors

The delivery of viral vectors and implantation of optic cannulas were conducted through stereotaxic surgeries. Mice were anesthetized with a ketamine/xylazine cocktail (100□mg/kg and 10□mg/kg, respectively, intraperitoneal), and their heads were affixed to a stereotaxic apparatus in absence of the pedal reflex. Viral vectors were delivered through a 0.5□µL syringe (Neuros Model 7000.5 KH, point style 3; Hamilton, Reno, NV, USA) mounted on a motorized stereotaxic injector (Quintessential Stereotaxic Injector; Stoelting, Wood Dale, IL, USA) at a rate of 30□nL/min. Viral preparations were titered at ∼10^12^ particles/mL. Volumes and coordinates for viral injections were as follows: 50∼75 nl/side, anteroposterior (AP) +1.25 mm, mediolateral (ML) ±0.2 mm, dorsoventral (DV) -4.95 mm for the BF; 50∼75 nl/side, AP -3.1 mm, ML ±0.3 mm, DV -4.6 mm for the VTA; 100 nl/side, AP +1.4 mm, ML -1.0 mm, DV -4.5 mm for the NAc. For optogenetic experiments, customized fiber optic cannulas [Ø1.25-mm stainless ferrule, Ø200-μm core, 0.39 numerical aperture (NA), 4.7 mm; Inper, Zhejiang, China] were implanted to target the VTA (AP -3.1 mm, ML -0.3 mm, DV -4.4 mm). For fiber photometry experiments, customized wide-aperture fiber optic cannulas (Ø1.25-mm stainless ferrule, Ø400-μm core, 0.66 NA, 5.0 mm; Doric Lenses, Quebec, QC, Canada) were implanted in the BF (AP +1.2 mm, ML -0.2 mm, DV -4.7 mm), the VTA (10°, AP -3.1 mm, ML -0.3 mm, DV -4.7 mm) and the NAc (10°, AP +1.4 mm, ML -1.7 mm, DV -4.2 mm). For AP5 and CNQX in vivo infusions, an optofluid cannula with interchangeable injectors (M3, Ø200-μm core, 0.37 NA, 4 mm guiding tube, 4.7 mm fiber, 4.5 mm injector; Doric Lenses, Quebec, QC, Canada) was implanted in the VTA. For all cannula implants, fibers were fixed to the skull with glue and dental cement (Stoelting 51458; Stoelting co., IL, USA). For the following three days of neurosurgery, mice were treated with Caprofen (Rimadyl; Zoetis Inc, MI, USA; i.p., 5mg/kg) for pain relief.

**Table:** Viral vectors information.

**Table: Viral vectors information**
| Viral constructs | Sources |
| --- | --- |
| AAV5-Ef1a-DIO-hChR2(H134R)-EYFP | Addgene #20298 |
| AAV-DJ8-hSyn-Con/Fon-EYFP-WPRE | UNC GTC Vector Core |
| AAVrg-Ef1a-mCherry-IRES-Flp | Addgene #55634 |
| AAV-DJ8-CAG-Flex-WGA-Egfp | Canadian Neurophotronics Platform |
| AAV5-hSyn-DIO-mCherry | Addgene #50459 |
| AAV1-Ef1a-Flp-DOG-NW | Addgene #75469 |
| AAV8-Ef1a-Con/Fon-mCherry | Addgene #137132 |
| AAV-DJ8-CAG-DIO-EGFP-TTC | Canadian Neurophotronics Platform |
| AAV-DJ8-Ef1a-fDIO-EYFP | UNC GTC Vector Core |
| AAV9-hSyn-FLEX-GCaMP6m | Addgene #100838 |
| AAV9-hSyn-DIO-hm4D(Gi)-mcherry | Addgene #44362 |
| AAV9-hSyn-DIO-hM3D(Gq)-mCherry | Addgene #44361 |
| AAV-DJ8-Ef1a-DIO-NaChBac-EGFP | Baylor Gene Vector Core |
| AAV5-Ef1a-DIO-EYFP | Addgene #27056 |
| AAV9-hSyn-g-GRAB-DA3m (gDA3m) | WZ Biosciences |
| AAV5-Ef1a-fDIO-mCherry | Addgene #114471 |

### Fiber photometry

All fiber photometry recordings were conducted using a Doric Lenses setup, with a light-emitting diode (LED) driver controlling three connectorized LEDs (405, 465) routed through a 5-port Fluorescence MiniCubes to deliver excitation light for isosbestic and calcium or dopamine-dependent signals to the implanted optic fiber simultaneously. Emitted light was collected by the same fiber and focused onto separate photoreceivers (Newport 2151) based on the wavelength pf fluorescent light. A Doric Fiber Console controlled by the Doric Studios (V6.1.4.0) was used to control the LEDs and demodulate the collected signals based on the wavelength.

For dual-cannula recordings in the BF and the VTA, we used the following experimental procedure. Animals were habituated in the recording cage for 10 minutes with the optic fiber connected. After habituation, signals and mice behaviors were recorded for 10 minutes. The first stimulus was delivered 2 minutes post recording initiation and each stimulus was given at least 1 minute apart. The water spray, air puff and object drop were presented to mice on different dates, at least three days apart.

For single-cannula recordings in the NAc, we used the following experimental procedure. Animals were habituated in the recording cage for 10 minutes with the optic fiber connected. After habituation, signals and mice behaviors were recorded for 10 minutes. The first stimulus (water spray or photostimulation (20HZ, 20ms, 10s or 20s) was delivered 2 minutes post recording initiation and each stimulus was given at least 1 minute apart. For feeding assay, chow diet-fed mice which had been pre-exposed to HFD were given the free access to HFD. The mice were habituated in the recording cage for 10 minutes with the presence of HFD pellets. For feeding assay with photostimulation, mice were also exposed to and given the free access to HFD pellets. Yet, during 10-min recording epochs, once the mice engaged with feeding, the photostimulation (20 HZ, 20 MS, 10 s) was applied to a separate optic cannula implanted in the VTA.

### *In vivo* photostimulation

Behavioral measurements were performed during the light cycle of the day at least four weeks post-surgery. An integrated rotary joint patch cable (Doric Lenses, Quebec, QC, Canada) was used to connect the ferrule of the implanted optic fiber to the 473-nm diode-pumped solid-state laser (Opto Engine LLC, Midvale, UT, USA). Light pulses (20 HZ, 20 MS) were controlled by Master-8 pulse stimulator (A.M.P.I., Jerusalem, Israel). Mice were placed in the sanitized Phenotyper cages (Noldus, Wageningen, Netherlands) with a camera real-time recording on top of the cage. After each test, cages were cleaned and sanitized with 75% ethanol. Before behavioral tests, mice were acclimated in the behavior room for at least 30 mins.

### Real-time place avoidance (RTPA)

The Noldus Phenotyper chamber was divided into two halves, one of which was paired with photostimulation. Mice were placed in the laser-off side in the beginning and were free to roam in the enclosure. The EthoVision XT software (version 15.0; Noldus) was triggered to collect tracking data for 10 minutes when mice entered the side paired with the photostimulation and the light pulses were applied. RTPA was repeated with the side paired with photostimulation counterbalanced after a week.

### Fasting refeeding assay

Mice were fasted overnight (16∼18 hours). Chow diet pellets were placed in a petri dish in a corner of Phenotyper cages. Mice were put in the center of cage and were free to roam in the enclosure. Once the mice entered the food corner and engaged in feeding, the Ethovision XT software was triggered to collect tracking data and apply laser pulses for 10 minutes. The weight of pellets consumed during the 10 minutes experiment episode were recorded. To deliver the glutamate receptor antagonists, the optic fiber cannula was removed from the guiding cannula and the injector was inserted. A syringe (0.5 µL, Model 7000.5KH, 25 ga, 2.75 in) joined with a plastic tube (RenaSil Silicone Rubber Tubing, .025 OD X .012 ID; Braintree Scientific, INC, Braintree, MA, USA) was used to deliver 50□nL (61□mM D-AP5 solution in saline)□+□50□nL (24□mM DNQX solution in 25–30% DMSO) cocktail. After 5 minutes incubation, the injector was removed, and the optic cannula was re-inserted, followed by feeding assays as described above.

### Laser on-off open field test

Mice were put in the center of Phenotyper cages and were acclimated for three minutes before the software tracking. Mice were free to roam in the enclosure and underwent 15-min trials consisting of three consecutive 5-min epochs (pre-laser, laser-on, and post-laser). During pre-laser and post-laser epochs, the laser was turned off. During the laser-on period, the laser (20 HZ, 20 MS) was applied. The total distance travelled, and the time spent in different zones were collected.

### Chemogenetics

Mice that received Gi or Gq viral vectors and control mice were treated with either saline or *Clozapine-N-oxide* (1 mg/kg, i.p.). Behavioral tests were conducted 30 mins post injections. For each cohort, half of mice were injected with saline and half of the mice were injected with CNO. After one week, experiments were repeated mice with saline and CNO injections in a counterbalance fashion. For metabolic cages recordings, mice were acclimated in metabolic cages for 2 days before application of saline/CNO at ZT 6.

### Singly housed fasting refeeding assay

Mice were singly housed and overnight fasted (16∼18 hours) before the assay. Mice were provided with standard chow pellets 30 mins after saline/CNO application. Food pellets were weighed at 0h, 0.5h, 1h, 2h, 4h, and 6h after mice accessed the food pellets.

### Operant conditioning

Mice were singly housed and kept with a calorie-restricted diet that they maintained 80∼85% of their original body weight for a week prior to training. Mice were trained daily for 30 mins in Nose Poke chambers (Nose Poke chambers, MED-307W-B2, Med Associates Inc, Fairfax, VT) on gradually-increasing fixed ratios (FR) of 1, 5, and 15. FR was increased whenever mice were able to acquire 25 pellets within 30 mins. At the test day, mice from both groups were injected with CNO and tested for positive reinforcements. Three days later, mice were tested again with saline injections.

### LDT and OFT

In LDT and OFT experiments, mouse movement was recorded with cameras mounted on top of the maze, the box and the phenotyper and tracked for 10 minutes using Ethovision XT software. The duration and frequency to access open arms, the light side chamber and the center of chamber were collected. For individual mice that went through consecutive tests in weeks, the order was LDT and then OFT.

## Physiology assessment

### Metabolic cages

Mice were individually housed in chambers of Columbus Instruments Comprehensive Lab Animal Monitoring System (Columbus Instruments, Columbus, Ohio, USA) for chemogentic experiments or the PhenoMaster cages (TSE systems, Chesterfield, Missouri, USA) for long-term body weight monitoring experiments. Mice were given ad libitum access to a normal chow diet and water. Food intake, O2 consumption, and locomotion activity levels were measured using indirect calorimetry continuously at different time points. Data was averaged through different time points in light and dark cycles, respectively, for comparison. The data from the first day and the last day was removed.

### Body weight and food intake measurement

Long-term body weights were collected weekly for 8 weeks post-surgery under normal chow diet and for another 8 weeks under high-fat diet (Research Diets D12492; 20% protein, 60% fat, 20% carbohydrate, 5.21 kcal/g). Ad libitum food intake was measured at the end of metabolic cages assessment and mice were singly housed. Daily food intake was recorded and averaged for three days.

## Post-hoc analysis

### Perfusion and tissue dissection

To harvest brain tissues, mice were anesthetized with ketamine/xylazine (150□mg/kg and 15□mg/kg, respectively). After loss of the pedal reflex, mice were transcardially perfused with 15 ml of saline and 15 ml of 10% buffered formalin (In Vivo Perfusion System IV-140, Braintree Scientific Inc.). The brains were then collected and stored in 10% buffered formalin overnight at room temperature, and then, the brain was switched to 30% sucrose in PBS for overnight. Brains were sectioned into 30 μM coronal slices on a frozen sliding microtome and stored in 0.1% NaN3 in PBS at 4 °C.

### Immunohistochemistry (IHC)

For IHC, sectioned slices were rinsed with 0.3% Triton X-100 in phosphate-buffered saline (PBS) for 5 mins three times and were blocked in 0.3% Triton X-100 in PBS with 10% donkey serum at room temperature (RT) for 1 hour. The slices were incubated in a primary antibody solution (primary antibody, 5% donkey serum, and 0.3% Triton X-100 in PBS) overnight at 4°C. The following primary antibodies were used: c-Fos rabbit monoclonal antibody (mAb) (9F6) (1:1000, #2250; Cell Signaling Technology), tyrosine hydroxylase rabbit polyclonal antibody (1:1000, ab112; abcam). For secondary antibody treatment, slices were rinsed with 0.3% Triton X-100 IN PBS for 5 mins for three times and incubated in the secondary antibody solution (Alexa Fluor 647–conjugated AffiniPure Donkey (H+L) anti-rabbit immunoglobulin G (Jackson ImmunoResearch), 1:400, 10% donkey serum, and 0.3% Triton X-100 in PBS) for 2 hours at RT. The floating slices were mounted onto microscope slides and coverslipped with Fluoromount (Diagnostic BioSystems Inc., Sigma-Aldrich). A confocal microscope was used to image the slices at different resolutions (Leica TCS SP5, Leica Microsystems, Wetzlar, Germany). Mice with offsite injections and cannula implants were removed from the study. The fluorescence-positive neurons were recognized and quantified in the software QuPath 0.3.2 (https://qupath.github.io).

## Statistics

GraphPad Prism 9.4.0 (GraphPad Software, Inc., La Jolla, CA, USA) was used for all statistical analyses and construction parts of the figures. For fiber photometry data, raw data from single trials was first processed through pMat application (https://github.com/djamesbarker/pMAT)^76^. Individual traces were aligned to zero at stimulus onset and averaged to be visualized in MATLAB R2022b (student version). In the heatmaps, signals were normalized from 0 to 1 across each trial. The unpaired two-tailed Student’s t-test was used for single-variable comparisons. Two-way repeated ANOVA followed by Sidak’s multiple comparisons were used for repeated measurements in group comparisons. Two-way non-repeated ANOVA followed by Sidak’s multiple comparisons was used for group comparisons. Error bars in graphs were represented as mean□± s.e.m. p < 0.05 was considered significant. p* < 0.05, p** < 0.01, p*** < 0.005, p**** < 0.0001. The sample sizes were chosen based on previously published work. All tests met assumptions for normal distribution, with similar variance between groups that were statistically compared. N values represent the final number of animals used in experiments following genotype verification and post-hoc validation of injection sites/cannula implantations.

## Acknowledgement

We acknowledge Dr. Yulong Li for providing the gDA3m vector. This study was supported by the NIH R01 DK114279, NIH R01 DK120858 and R01DK136284 (QT), R01DK109934 and DOD W81XWH-19-1-0429 (QT and BRA). QT is the holder of the Cullen Chair in Molecular Medicine at McGovern Medical School. JC is the awardee of Russell and Diana Hawkins Family Foundation Discovery Fellowship.

## Author contributions

JC conducted the major part of research with help from YX, YJ, and ZJ. JO-G, BRA, YZ, and YL provided essential reagents; QT and BRA conceived and designed the experiments, and wrote the manuscript with significant inputs from all authors.

