## Supplementary figures for "An Excitatory Projection from the Basal Forebrain to the Ventral Tegmental Area Underlying Anorexia-like Phenotypes"

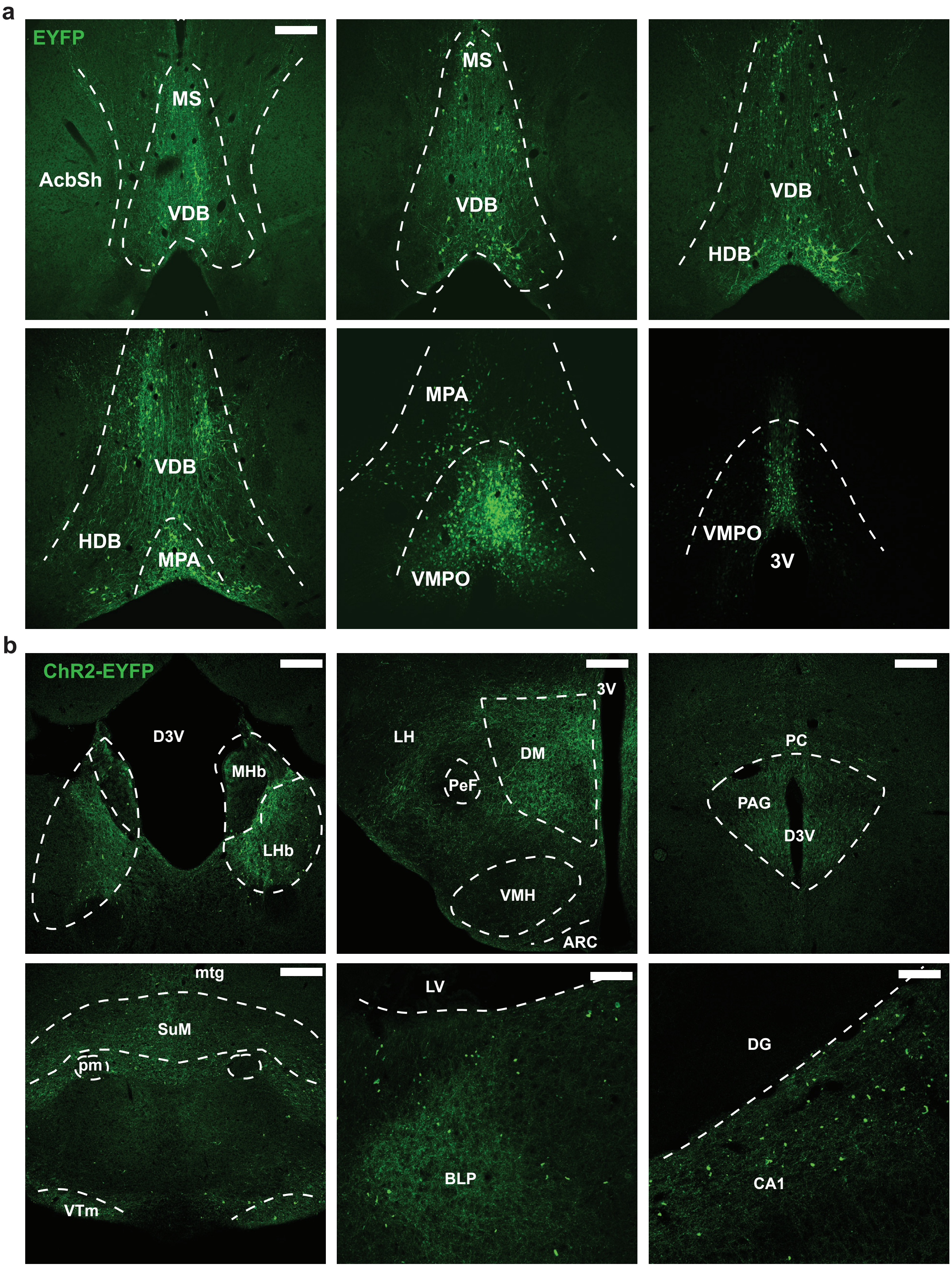


**Supplement Figure 1: BF glu+ neurons project to other brain regions in addition to the VTA**. a, Representative coronal sections of the BF with Cre-dependent GFP expressions, from anterior to posterior. AAV5-hSyn-DIO-EYFP was injected to the BF of Vglut2-iCre mice unilaterally. AcbSh: the nucleus accumbens shell. MPA: media preoptic area. VMPO: ventral media preoptic area. b, ChR2 positive neuronal fibers from the BF Vglut2 neurons in downstream regions. AAV5-hSyn-DIO-ChR2-EYFP was injected to the BF unilaterally. D3V: dorsal third ventricle. PeF: perifornical nucleus. DMH: dorsal medial hypothalamus. VMH: ventral medial hypothalamus. ARC: arcuate nucleus. pc: posterior commissure. PAG: periaqueductal gray. mtg: mammillotegmental tract. SuM: supramammillary nucleus. pm: principle mamaliary nucleus.  VTm: ventral tuberomammillary nucleus. LV: lateral ventricle. BLP: basolateral amygdala nucleus. DG: dentate gyrus. CA1: hippocampal CA1 region. Scale bar = 200 μm.

a

| Bregma (mm) | -3.08 | -3.16 | -3.28 | -3.4 |
|---|---|---|---|---|
| Mouse 1 | 93.5294 | 42.4821 | 50.3467 | 76.3689 |
| Mouse 2 | 68.8092 | 74.6324 | 18.1556 | 45.403 |
| Mouse 3 | 54.0779 | 31.6877 | 23.026 | 4.13462 |
| Mouse 4 | 93.6893 | 95.5882 | 54.3326 | 55.5932 |

b

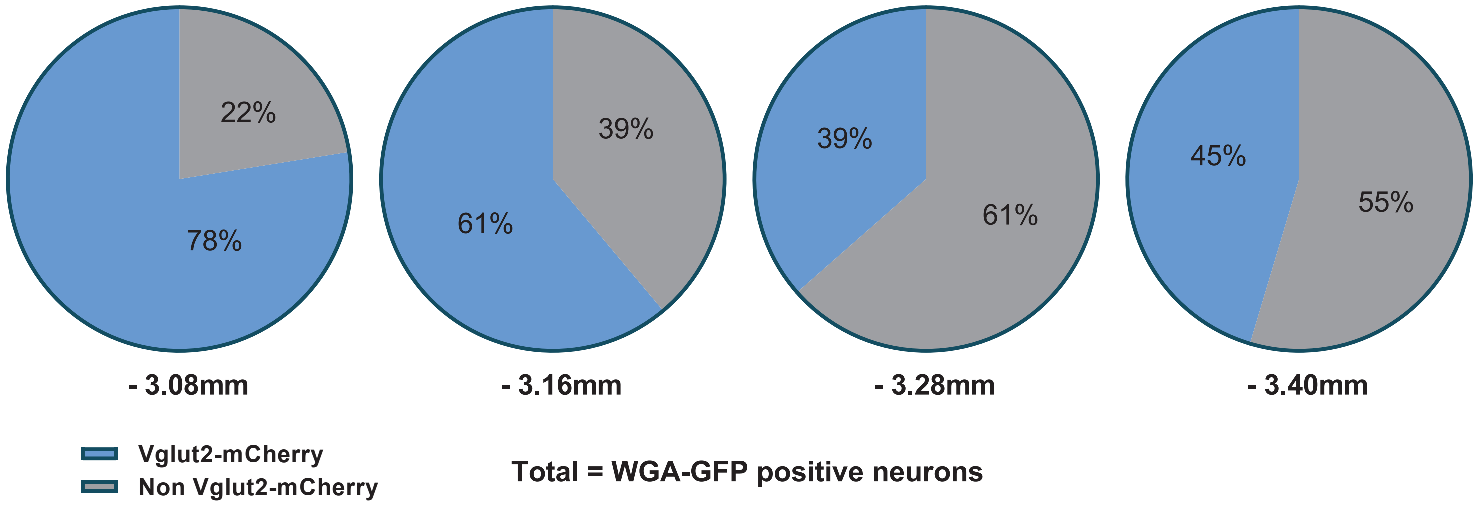


**Supplement Figure 2: Qualitative results of proportions of glu+ neurons in VTA neurons that form synaptic connections with BF glu+ neurons.** Conditional AAV-DJ8-Flex-WGA-EGFP viral particles to the BF of Vglut2-ires-Cre mice. WGA or EGFP in the VTA labelled VTA neurons that form synapse connections with BF glu+ neurons. AAV5-hSyn-DIO-mCherry viral particles were delivered to label VTA glu+ neurons. a, Quantification results of proportions of glu+ neurons (mCherry positive) in VTA neurons that form synaptic connections with BF glu+ neurons (EGFP positive) in individual animals. b, Pie plot of summarized quantification.

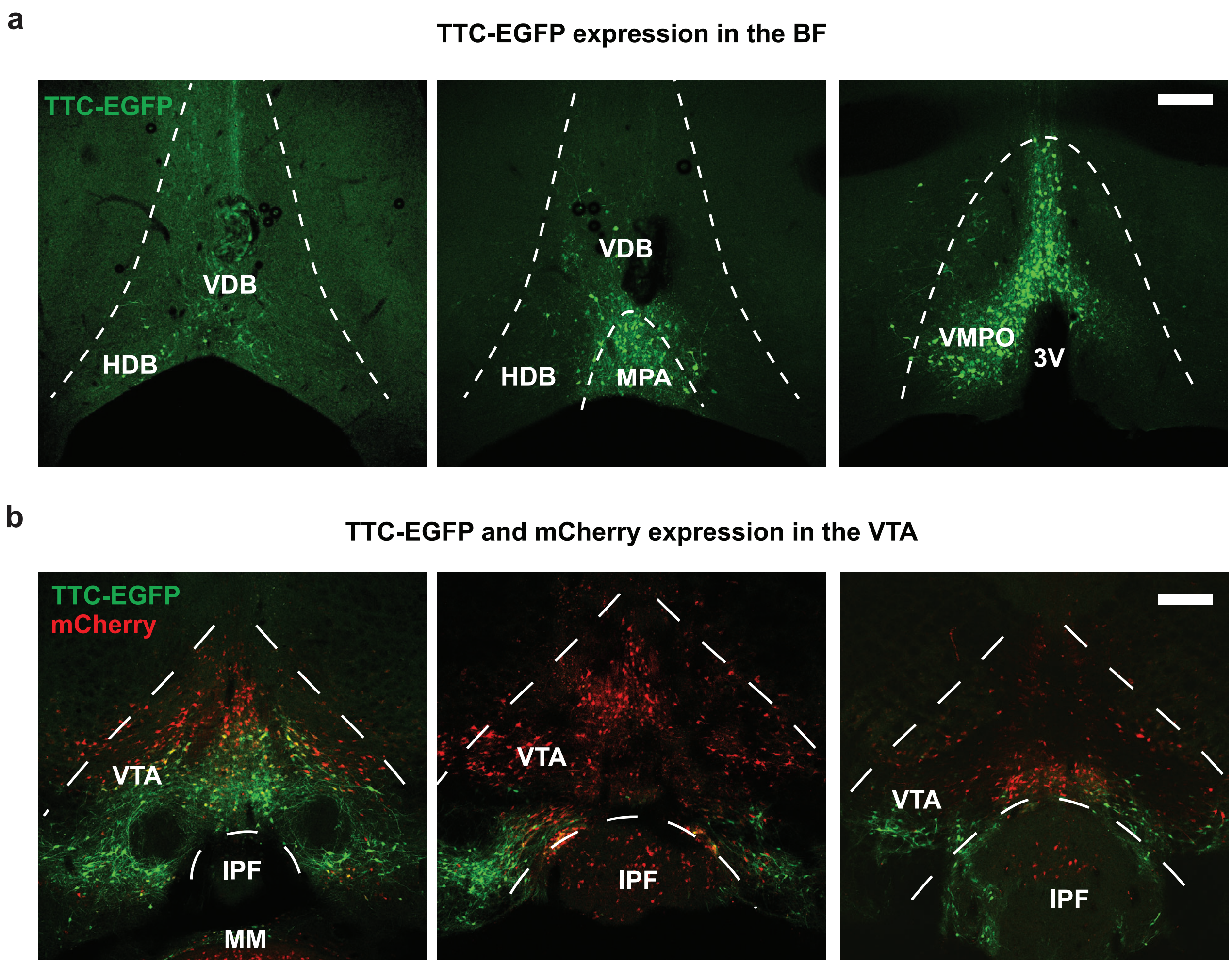


**Supplement Figure 3: VTA non-glu+ neurons project to BF$^{glu+}$ neurons.** AAVDJ8-CAG-DIO-TTC-EGFP was delivered in the BF, and AAV5-hSyn-DIO-mCherry was delivered in the VTA. The signal of TTC-EGFP was amplified with co-delivery of AAV1-DOG-Flp and AAVDJ8-EF1a-fDIO-EYFP in the VTA. a, Injection patterns of TTC-EGFP in the BF. b, Virus expression patterns in the BF. EGFP labelled neurons that send projections to the BF. The majority of EGFP-labeled neurons are not mCherry or Vglut2-ires-Cre positive. Scale bar = 200 μm.

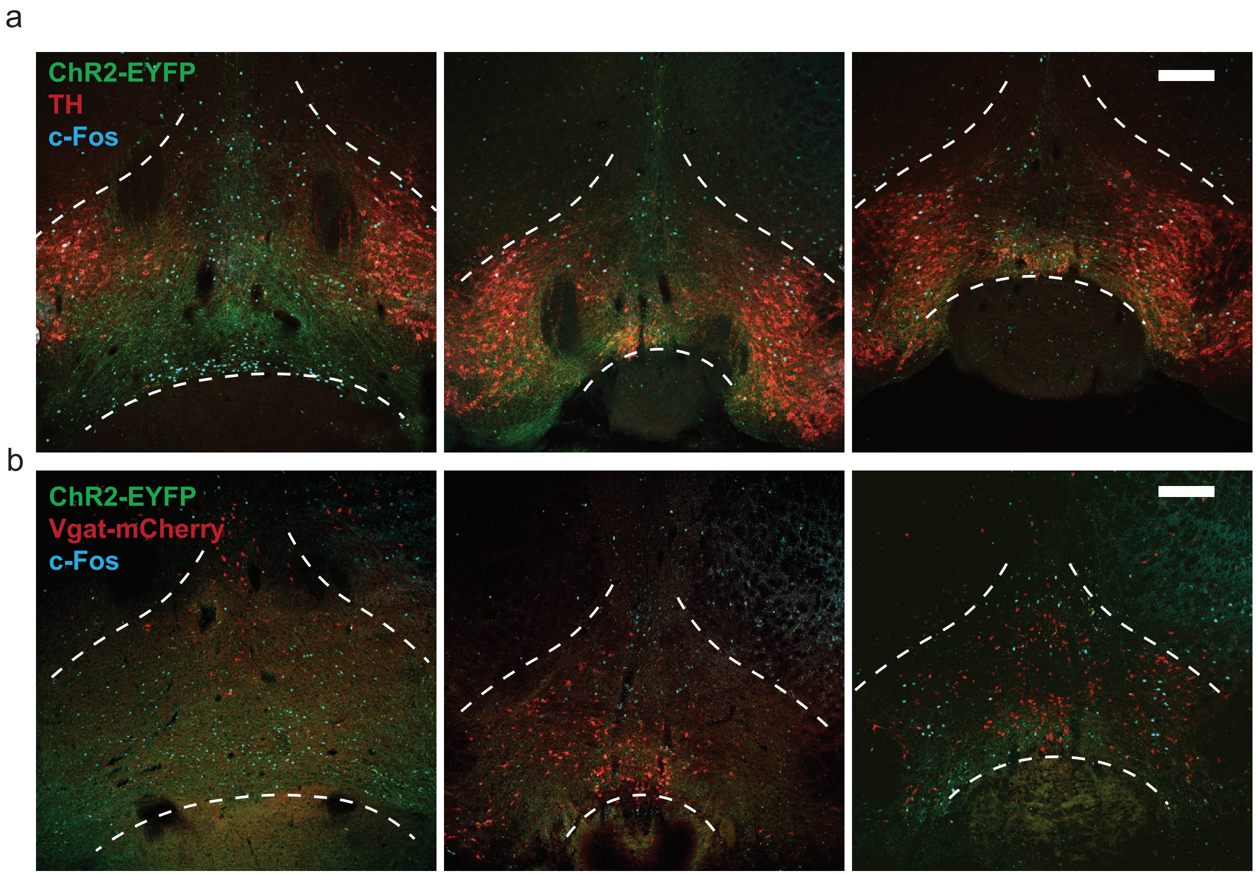


**Supplement Figure 4: c-Fos expression induced by activation of BF$^{glu+}$ → VTA displayed minimal co-localization with VTA DA and GABA neurons.** AAV5-EF1a-DIO-ChR2-EYFP was delivered in the BF of Vglut2-ires-Cre mice and the optical cannula was implanted in the VTA. The laser (λ = 473 nm) stimulation intensity is 20HZ, 20ms for 10 minutes. Tyrosine hydroxylase (TH) immunostaining was used to visualize VTA DA neurons. To visualize VTA GABA neurons, Vglut2-ires-Cre mice were crossed with Vgat-ires-Flp mice and AAV5-EF1a-fDIO-mCherrry vector was injected to the VTA. a, Anterior to posterior sections of the VTA showing TH neurons displayed limited c-Fos expression. b, Anterior to posterior sections of the VTA showing Vgat neurons displayed limited c-Fos expression. Scale bar = 200 µm.

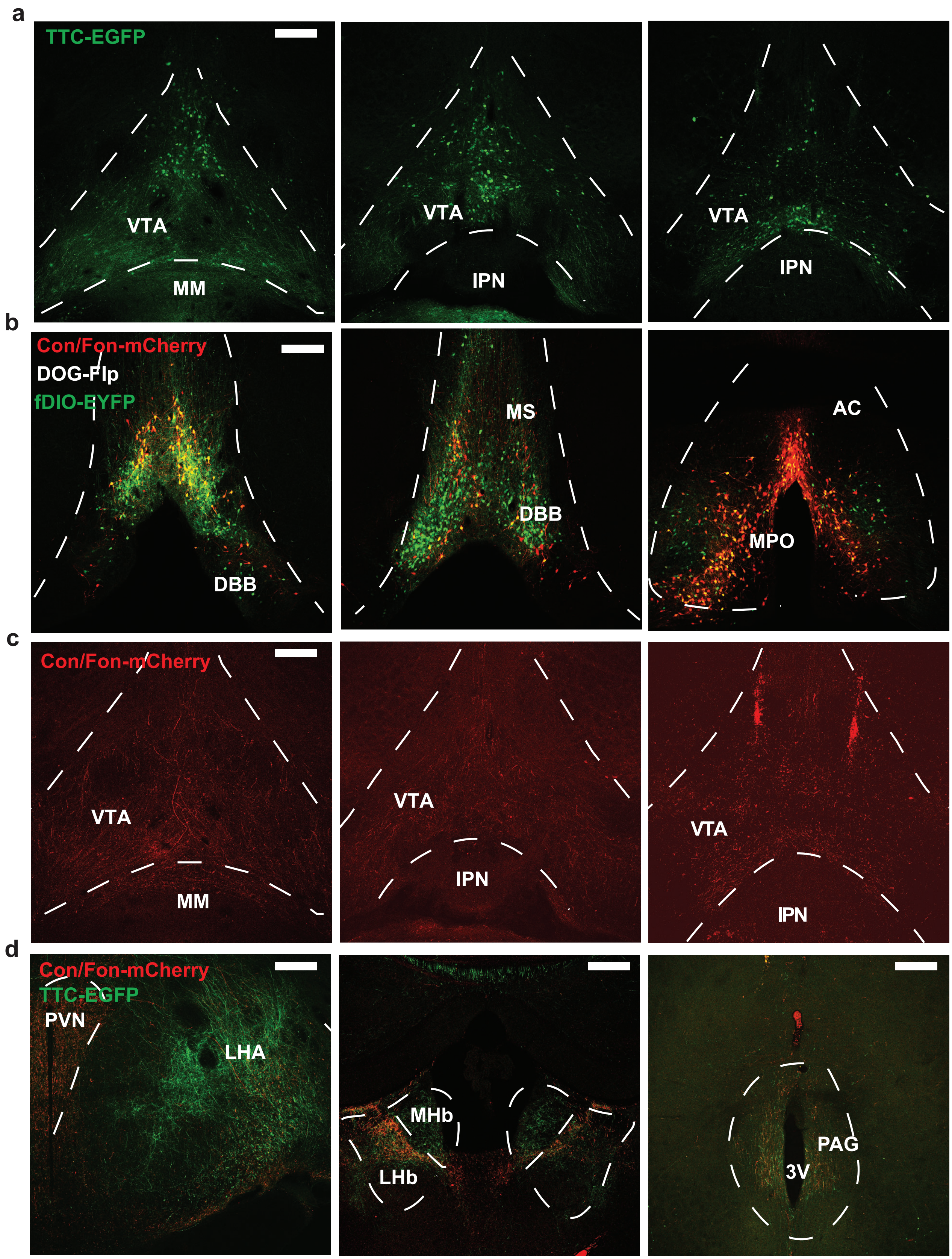


**Supplement Figure 5: VTA$^{glu+}$ neurons projecting BF$^{glu+}$ neurons send collateral projections to other brain regions in addition to the VTA.** AAVDJ8-CAG-DIO-TTC-EGFP was delivered in the VTA , and AAV8-EF1a-Con/Fon-mCherry was delivered in the BF. The signals of TTC-EGFPwere amplified with a co-delivery of AAV1-DOG-Flp and AAVDJ8-EF1a-fDIO-EYFP in the BF. a, Injection patterns of TTC-EGFP in the VTA. b, Virus expression patterns in the BF. Con/Fon-mCherry labeled BF$^{vglu+}$ neurons that send projections to the VTA glu+ neurons. c, Con/Fon-mCherry positive neuronal fibers from the BF in the VTA. d, Collateral projections from VTA$^{glu+}$ neurons projecting BF$^{glu+}$ in the hypothal amus, LHb, and the PAG. Scale bar = 200 μm.

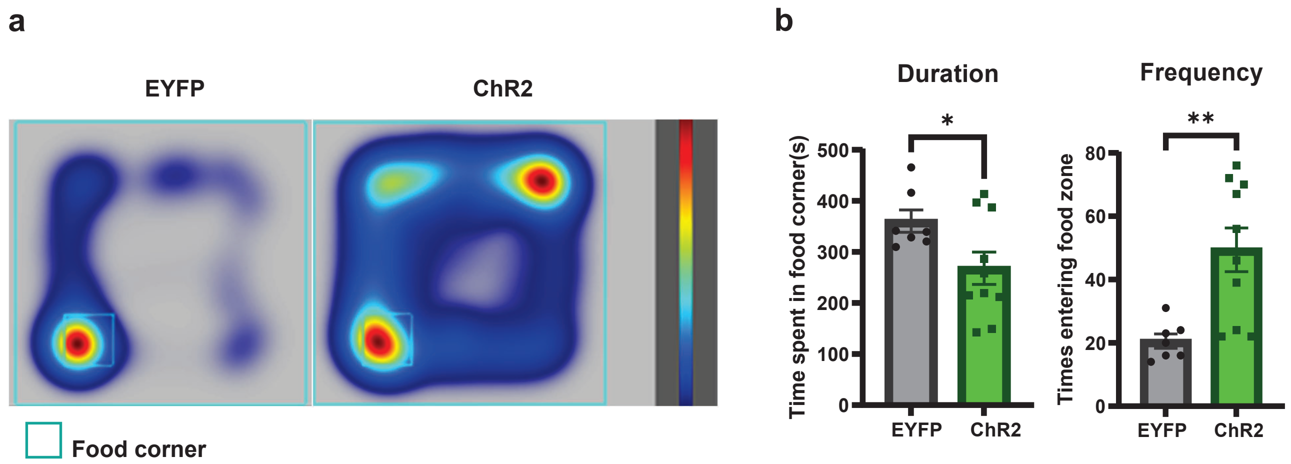


**Supplement Figure 6: Activation of BF$^{glu+}$ → VTA$^{glu+}$ projections reduced food consumption time.** AAV5-EF1a-DIO-ChR2-EGFP was delivered in the BF of Vglut2-ires-Cre mice and the optical cannula was implanted in the VTA. The laser (λ = 473 nm) stimulation intensity is 20HZ, 20ms for 10 minutes. Mice were able to move freely in the chamber and had free access to the food corner. a, Heatmap of mice movements in the chamber for 10 mins with photostimulation on. b, Qualitativeresults of duration and frequency of mice enter the food corner. Unpaired t-tests: duration, P = 0.0459; frequency: P = 0.0041.

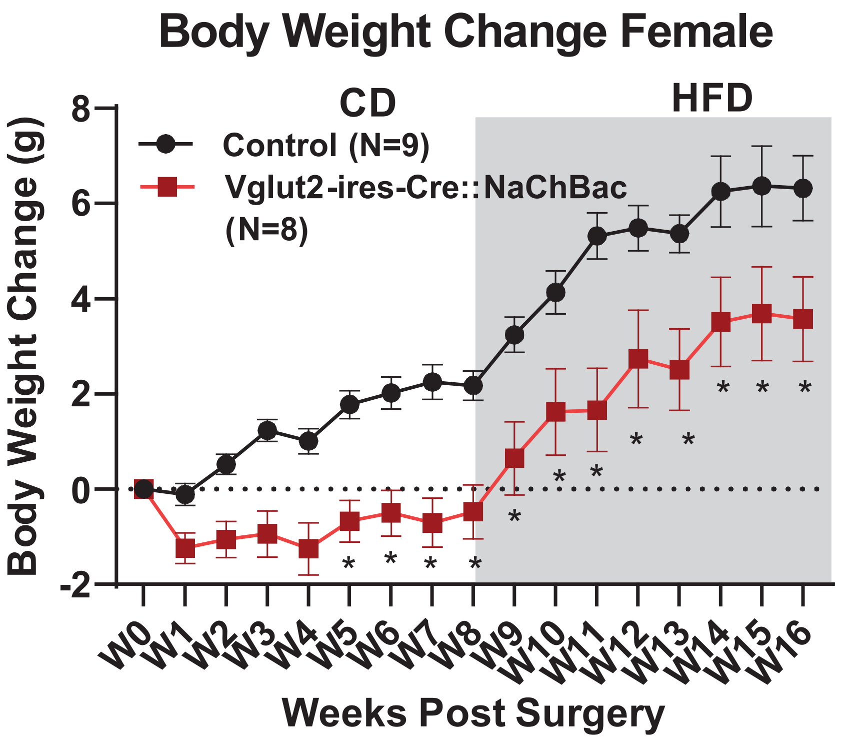


**Supplement Figure 7: Chronic activation of VTA glu+ neurons r educed body weight in female mice.** AAV-DJ8-EF1a-DIO-NaChB acEYFP and AAV5-EF1a-DIO-EYFP viral vectors were delivered to the VTA of Vglut2-ires-Cre mice. The body weight change of 16 weeks after surgery. From 0 to 8 weeks post-surgery, mice were fed with chow diet. From 9 to 16 weeks post-surgery, mice were fed with high fat diet (HFD). Two-way repeated ANOVA followed by Sidak m ultiple comparisons test: $F (16, 240) = 2.351$, $P = 0.0029$

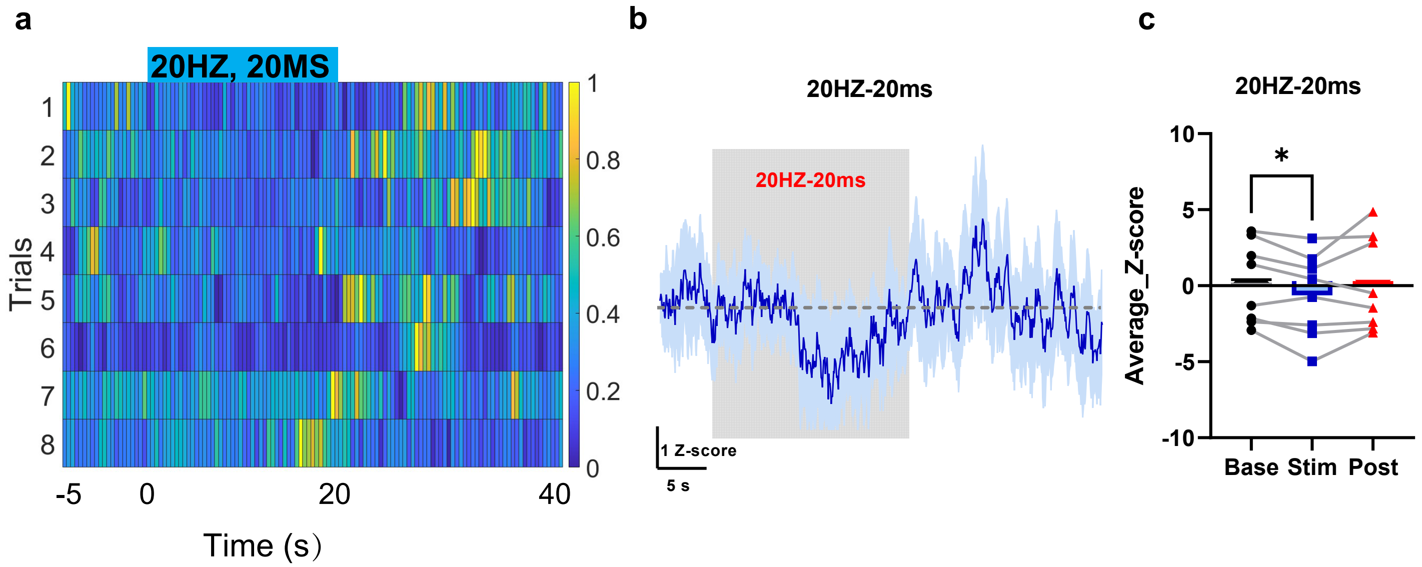


**Extended Fig 8: DA changes were inhibited upon activation of the BF-->VTA circuit.** AAV9-hSyn-GRABDA3.3 viral vectors were injected and an optic cannula was implanted in the NAc. AAV5-EF1a-DIO-ChR2-EYFP was expressed in the BF of Vglut2-ires-Cre mice and an optic cannula was implanted in the VTA. a, Heatmap presented DA Z-score signals of individual trials, signals were re-scaled from 0 to 1 across each row. The indigo box suggested the application of laser. b, Traces represented average DA Z-score signals. The shade represented +/- SEM. Time = 0 was the onset of laser. c, Averaged Z-scores from baseline, laser application, and 5 seconds post laser application. Paired t-test: Stim vs. Base, P = 0.023; Post vs. Stim, P = 0.1886.
